# Human memory CD4^+^ T-cells recognize *Mycobacterium tuberculosis*-infected macrophages amid broader pathogen-specific responses

**DOI:** 10.1101/2025.02.23.639515

**Authors:** Volodymyr Stetsenko, Daniel P. Gail, Scott Reba, Vinicius G. Suzart, Avinaash K. Sandhu, Alessandro Sette, Mohammad Haj Dezfulian, Cecilia S. Lindestam Arlehamn, Stephen M. Carpenter

## Abstract

Recognition of macrophages infected with *Mycobacterium tuberculosis* (Mtb) is essential for CD4^+^ T cells to prevent tuberculosis (TB). Yet not all antigen-specific T cells recognize infected macrophages in human and murine models. Using monocyte-derived macrophages (MDMs) and autologous memory CD4^+^ T cells from individuals with latent Mtb infection (LTBI), we quantify T cell activation in response to infected macrophages. T cell antigen receptor (TCR) sequencing revealed >70% of unique and >90% of total Mtb-specific TCR clonotypes in stable LTBI are linked to recognition of infected macrophages, while a subset required exogenous antigen exposure, suggesting incomplete recognition. Clonotypes specific for multiple Mtb antigens and other pathogens were identified, indicating Mtb-specific and non-specific activation. Single-cell transcriptomics demonstrates Mtb-specific T cells express signature effector functions dominated by IFNγ, TNF, IL-2, and GM-CSF or chemokine production and signaling. We propose TB vaccines that elicit T cells capable of recognizing infected macrophages and expressing these canonical effector functions will offer protection against TB.

## Introduction

*Mycobacterium tuberculosis* (Mtb), the bacterium that causes tuberculosis (TB), continues to cause more deaths than any other infectious disease (Bagcchi, 2023). There is an urgent need for a vaccine that prevents active TB and reduces Mtb transmission. However, discovering the immune cells with the greatest potential for protection through vaccination has proven challenging. CD4⁺ T cells are a major focus of TB vaccine development due to their central and multifaceted role in the immune response to Mtb infection (Bell and Noursadeghi, 2018; Mogues et al., 2001; Bromley et al., 2024). Protection by CD4^+^ T cells was shown to depend on the direct recognition of peptide-major histocompatibility complexes (pMHC) expressed on infected antigen-presenting cells (APCs) by Mtb-specific T cells (Srivastava and Ernst, 2013). Yet evidence from the mouse model of TB demonstrates a lack of recognition of infected macrophages by a large fraction of Mtb-specific T cells (Yang et al., 2018; Patankar et al., 2020; Bold et al., 2011). We and others have found macrophage phenotype (Gail et al., 2023), antigen specificity (Yang et al., 2018), the context of T cell priming (Griffiths et al., 2016; Nyendak et al., 2016; Patankar et al., 2020), TCR avidity and dissociation rate (Gallegos et al., 2016; Nauerth et al., 2013), and mycobacterial virulence factors (Mwebaza et al., 2023) each contribute to T cell recognition of infected cells. For humans, the extent to which CD4^+^ T cell recognize infected macrophages is unexplored.

Upon infection, effector T cells are identified in the lungs only after priming occurs (Wolf et al., 2008; Reiley et al., 2008; Carpenter et al., 2017, 2016). T cell priming depends on the transport of live bacteria from the lungs by CCR2^+^ monocyte-derived cells, and on antigen presentation by conventional dendritic cells (cDCs) in mediastinal lymph nodes (MLNs) (Samstein et al., 2013; Wolf et al., 2008). Yet, it is macrophages in the lungs that serve as the niche cells for Mtb growth (Russell et al., 2025), making their recognition by T cells critical to protection. Differences in Mtb antigen presentation by DCs and infected macrophages in the lungs could explain the lack of recognition by a subset of T cells. Using mass spectroscopy, a recent study found only a fraction of the Mtb proteome to be presented by human monocyte-derived APCs after infection (Leddy et al., 2023). Since direct measurements of the Mtb peptides presented *in vivo* is impracticable in humans, indirect assessments using TCR sequencing of T cells activated in response to antigen-laden or infected APCs are critical tools.

Individuals who effectively control Mtb infection were found to possess TCRs specific for a range of Mtb antigens to facilitate efficient bacterial clearance (Musvosvi et al., 2023; Huang et al., 2020; Glanville et al., 2017). Conversely, individuals who fail to control infection, and ultimately progress to active TB, exhibit enrichment of alternate TCR motifs (Musvosvi et al., 2023), indicating a difference in antigen specific T cell responses. In murine and non-human primate (NHP) models, the effector functions expressed by protective CD4^+^ T cells in the lungs include canonical IFNγ, TNF, and IL-2 production (Lewinsohn et al., 2017), GM-CSF (Rothchild et al., 2017; Dis et al., 2022), cytotoxicity (Gideon et al., 2022), and, depending on the context, IL-17 or IL-10 production (Dijkman et al., 2019; Bromley et al., 2024). In humans, LTBI that is “stable” (i.e., without progressing to active TB) has been associated with protection from TB (Andrews et al., 2012). Increased memory CD4^+^ T cell responses also correlated with protection among individuals vaccinated with the M72/AS01_E_ candidate TB vaccine (Rodo et al., 2019). Despite promising efficacy with M72/AS01_E_ (Tait et al., 2019), a major challenge in TB vaccine development is the lack of a reference for protective T cell function. In contrast, other TB vaccine candidates have not yet shown protection (Tameris et al., 2013; Darrah et al., 2014). In one case, human T cell clones derived from recipients of the AERAS-402 vaccine lacked recognition for Mtb-infected cells despite responding to peptide antigens (Nyendak et al., 2016). These findings suggest the ability to recognize infected macrophages could serve as marker of protective T cells if linked to clinical protection from TB.

In this study, we sought to quantify the proportion of peripheral blood human memory CD4^+^ T cells that recognize Mtb-infected macrophages. Using *ex vivo* co-culture of primary human CD4^+^ T cells with autologous Mtb-infected macrophages, we tested the hypothesis that a subset of circulating memory CD4^+^ T cells from individuals with LTBI do not efficiently recognize Mtb-infected macrophages. Among 10 individuals with stable LTBI, the majority of the Mtb-specific CD4^+^ T cells became activated in response to infected macrophages, whereas a subset required exogenous antigen exposure, suggesting incomplete recognition. We used single-cell TCR sequencing (scTCRseq) to refine our estimates of CD4^+^ T cell recognition of infected macrophages and were surprised to identify viral pathogen-specific TCR clonotypes, suggesting bystander activation. Strikingly, by clustering TCRs using Grouping of Lymphocyte Interactions by Paratope Hotspots V2 (GLIPH2), several of our top GLIPH2 groups were equivalent to those recently published, containing TCRs annotated as specific for the CFP10, EspA, and mIHF antigens (Glanville et al., 2017; Musvosvi et al., 2018; Huang et al., 2020). Antigen screening of another top GLIPH2 group responsive to infected macrophages revealed a TCR specific for the EccE3 antigen. Using single-cell transcriptomics (scRNAseq), we distinguished the effector functions of memory CD4^+^ T cells enriched with either Mtb-specific or viral antigen-specific TCRs. The identification of TCR clonotypes and effector functions linked to CD4^+^ T cell recognition of infected macrophages will inform TB vaccine design.

## Results

### A subset of memory CD4^+^ T cells lack recognition for Mtb-infected macrophages

To determine the proportion which recognize Mtb-infected macrophages, we isolated memory (CD45RA^Lo^) CD4^+^ T cells and CD14^+^ monocytes from the peripheral blood of healthy individuals with “stable” LTBI. We enrolled 10 healthy adult volunteers with a history of exposure to TB and a positive IFNγ release assay (IGRA) or tuberculin skin test (TST) > 10mm obtained at least 3 years prior to participating (see Methods). After MDM differentiation and infection with Mtb strain H37Rv at a MOI of 4-5, we co-cultured infected MDMs with autologous memory CD4^+^ T cells (**Fig. 1A**). We previously found that infection with H37Rv at a MOI of 4-5 leads to infection of >75% and 95% of M1 and M2-like human macrophages, respectively (Gail et al., 2023). 16-18h later, activation-induced markers (AIMs) and IFNγ secretion were used to quantify the subset of memory CD4^+^ T cells activated in response to Mtb-infected macrophages, as performed previously (Gail et al., 2023, 2024; Reiss et al., 2017; Huang et al., 2020; Musvosvi et al., 2023). To identify Mtb-specific memory CD4^+^ T cells that lacked efficient recognition, we compared the proportion of T cells activated in response to infected macrophages with infected macrophages to which MTB300 megapool peptides (Arlehamn et al., 2016) or H37Rv whole-cell lysate (lysate) were added (**Fig. 1A**). Consistent with our hypothesis, greater numbers of memory CD4^+^ T cells co-expressed the AIMs CD69 and CD40L in response to infected macrophages treated with MTB300 (0.753 ± 0.074%; mean ± SEM, 10 participants) or lysate (0.899 ± 0.069%) compared to infection-only (0.606 ± 0.054%) (**Figs. 1B-1E**). Interestingly, most but not all participants’ CD4^+^ T cell responses increased with the addition of MTB300 (8/10 individuals) or lysate (9/10 individuals), suggesting response heterogeneity among individuals (**Figs. 1D and 1E**). The same trend was observed for IFNγ^+^ secretion by CD69^+^ CD4^+^ T cells in response to infected macrophages (0.326 ± 0.040%; mean ± SEM, 6 participants) + MTB300 (0.678 ± 0.176%) or + lysate treatment (0.863 ± 0.097%) (**Figs. 1B and 1F**), and for CD69 and CD25 co-expression (**Figs. S1A-C**). The reduction in T cell activation with α-MHC-II (HLA-DR/-DQ/-DP) mAb blockade indicates TCR-mediated activation (**Fig. 1B-D and Fig. S1B**). The increased proportion of memory CD4^+^ T cells activated when MTB300 was added to infected macrophages preferentially occurred among participants with LTBI while little change occurred for healthy control participants without LTBI and a negative IGRA (non-LTBI) (**Figs. 1E, 1G, and Fig. S1D**). Therefore, the increase in activated T cells observed when exogenous Mtb antigens were added suggested dominant but incomplete recognition of infected macrophages by Mtb-specific memory CD4^+^ T cells.

**Figure 1.**
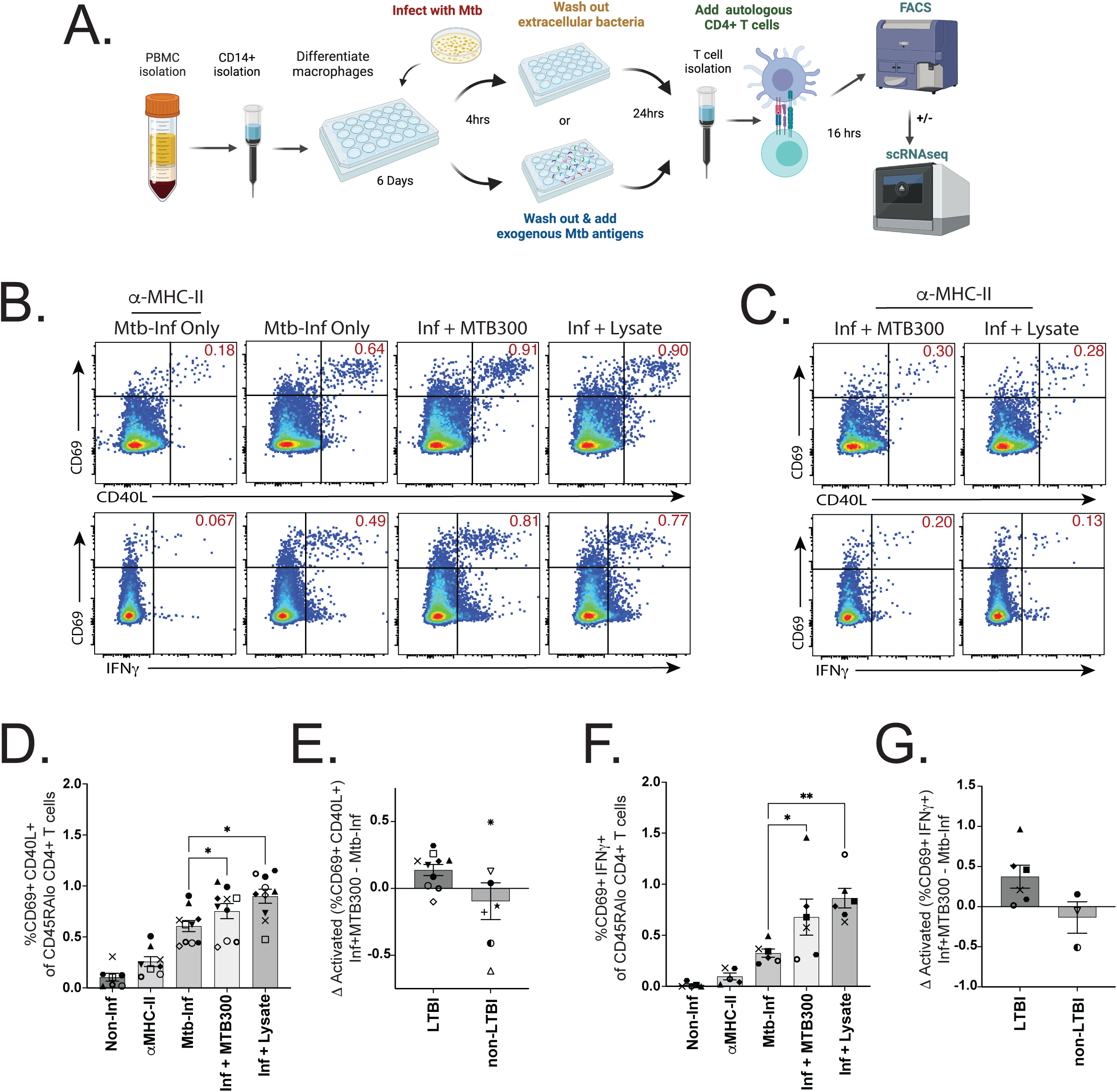
A subset of memory CD4^+^ T cells lack recognition for Mtb-infected macrophages. **(A)** Schematic of experimental workflow to co-culture infected macrophages with autologous memory CD4^+^ T cells for flow cytometry or sorting. Created in BioRender. Carpenter, S. (2025). **(B)** Representative flow plots comparing activation marker co-expression of CD69 with CD40L (top row) or IFNγ (bottom row), gated on CD45RA^Lo^ CD4^+^ T cells after 16-18 h co-culture with Mtb infected macrophages ± treatment with MTB300 or lysate, and **(C)** in the presence of α-MHC-II blocking antibodies. Data are representative of 10 (CD69 vs CD40L) and 6 (CD69 vs IFNγ) experiments and participants. **(D)** Summary bar graphs compare mean (± SEM) co-expression of CD69 and CD40L, and **(E)** the difference in activation when MTB300 is added to infected macrophages (10 LTBI and 7 non-LTBI). **(F)** Summary bar graphs compare mean (± SEM) CD69 and IFNγ co-expression, and **(G)** change in activation when MTB300 is added (6 LTBI and 3 non-LTBI). Each symbol represents the mean of 1-3 replicates from independent experiments. Statistical significance was determined by Wilcoxon matched pairs signed rank test. * p < 0.05, ** p < 0.01. See also Supplemental Figure 1.

### scTCRseq identifies clonotypes linked to recognition of infected macrophages

Although AIMs are useful for identifying antigen-specific T cells in response to peptide stimulation, additional inflammatory cytokine-mediated “bystander” activation can be elicited by infection or lysate treatment, leading to an overestimation of the Mtb-specific response(Reiss et al., 2017). Therefore, we performed αβTCR sequencing on CD4^+^ T cells activated in response to infected macrophages to refine our estimates of Mtb-infected macrophage recognition. Following co-culture with infected MDMs (± MTB300 or lysate), AIM+ memory CD4^+^ T cells were flow-sorted and high-throughput single-cell sequencing was performed. Among the top 50 CDR3β sequences from a representative LTBI and non-LTBI participants, clonally expanded CDR3β sequences, including ‘CASSPGTESNQPQHF’, were identified in LTBI (**Figs. 2A and 2B**). The ‘TESN’ motif in this TCR was previously linked to specificity for the Mtb antigen EspA_301-315_ in a South African cohort with LTBI (Huang et al., 2020). Greater TCR clonality was observed among individuals with LTBI, compared to non-LTBI controls as indicated by lower Shannon and Inverse Simpson indices (**Figs. 2C and 2D**). Among individuals with LTBI, a mean of ∼40% of unique TCRs were clonally expanded, compared to ∼20% for non-LTBI (**Fig. 2E, left panel**). Upon examining TCRs present in ≥ 3 or ≥ 4 copies, more expanded clonotypes were identified for LTBI vs. non-LTBI participants at the expense of fewer total TCRs (**Fig. 2E, middle and right panels**). Since 16-18h is not sufficient to elicit T cell proliferation *in vitro*, clonal T cell expansions were determined to have previously occurred *in vivo* in response to infection.

**Figure 2.**
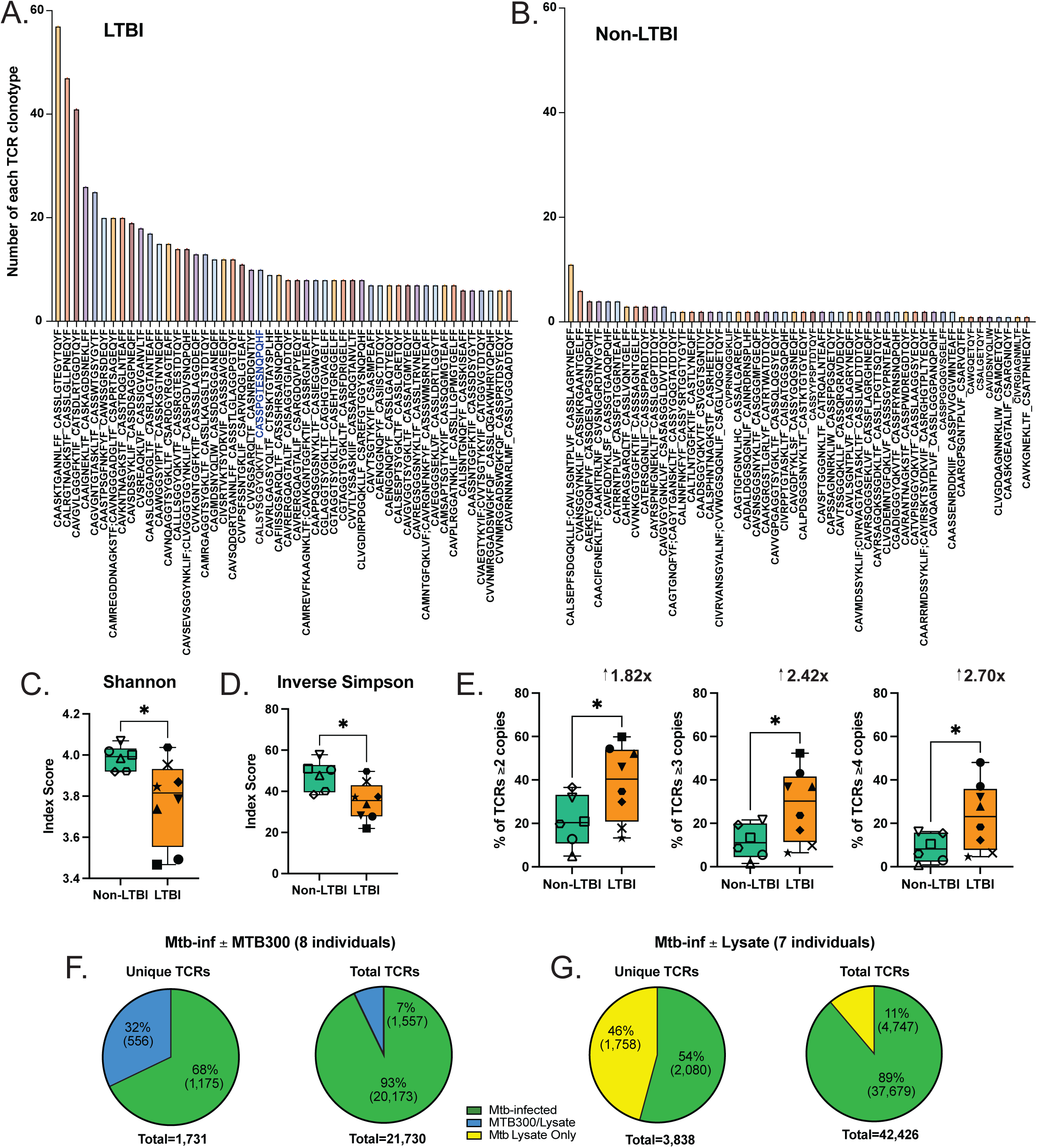
scTCRseq identifies clonotypes linked to recognition of infected macrophages. **(A)** Bar graphs of the top 50 TCR clonotypes and their frequencies from representatives of 8 LTBI and **(B)** 6 non-LTBI participants. Clonotypes are displayed as CDR3α_CDR3β sequences; some TCRs contained two CDR3α or β chains. Blue font highlights a CDR3β motif previously annotated as specific for EspA_301-315_. **(C)** Summary box plots of the average Shannon and **(D)** Inverse Simpson diversity index scores of 8 LTBI and 6 non-LTBI participants. Each symbol represents individual participant TCR repertoires. **(E)** Summary box plots of %TCR clonotypes present in ≥ 2, 3, or 4 copies with fold-differences listed above each graph. Statistical significance was determined by unpaired t test with Welch’s correction. **(F)** Pie charts combining TCRs from 7-8 participants, each, showing percent (and number) of unique TCRβ sequences linked to memory CD4^+^ T cell activation in response to infected macrophages (green) or after adding MTB300 (blue), or **(G)** lysate (yellow). Plots include all unique TCR clonotypes that meet each expansion threshold (≥2, 3, or 4 copies). **(H)** Pie charts for percent (and number) of total TCRs for each are also plotted. * p < 0.05. See also Supplemental Figure 2.

We next sought to create a condition where a set of infected macrophages also received exogenous treatment with MTB300 peptides, followed by co-culture with autologous memory CD4^+^ T cells. This enabled us to compare the TCR repertoires among responses to Mtb-infected macrophages ± treatment with MTB300 peptides. We found the majority of unique and total expanded TCR clonotypes were linked to recognition of infected macrophages, but a subset of 7-32% required exposure to MTB300 (**Fig. 2F**). Like MTB300, new TCR clonotypes were also identified when lysate was added to infected macrophages (**Fig. 2G**). Yet the majority of both unique and total TCRs were again linked to recognition of infected macrophages. Therefore, scTCRseq, and a focus on expanded TCR clonotypes, revealed dominant but incomplete recognition of infected macrophages by memory CD4^+^ T cells.

### GLIPH2 refines estimates of Mtb-specific CD4^+^ T cell recognition of infected macrophages

We next took 3 approaches to focus our analysis on clonotypes that are Mtb-specific. First, we stimulated whole PBMCs from the same individuals with either MTB300 or a combination of “control” viral and vaccine peptide megapools from CMV, EBV, *Bortadella pertussis*, tetanus toxoid (Dan et al., 2016; Antunes et al., 2017; Pro et al., 2015) and the SARS-CoV-2 spike protein. After 16-18h, we flow-sorted (**Fig. S2**) the AIM^+^ CD4^+^ T cells and performed scTCRseq, identifying TCR clonotypes that responded either to MTB300 or control peptides. Following peptide stimulations, we cross-referenced TCRs within each GLIPH2 group with the responses to infected macrophages and separated those activated by control peptides (**Fig. 3A**). Second, we used GLIPH2 (Huang et al., 2020; Musvosvi et al., 2023) to group the expanded TCRs (≥ 2 copies) from the combined list of sequences from all experimental conditions, including responses to infected macrophages ± MTB300, infected macrophages ± lysate, and peptide stimulations of PBMCs using either MTB300 or control peptide pools. After removing groups that contained TCRs linked to control peptide responses (**Fig. 3A**), we identified 107 GLIPH2 groups that met statistical criteria for CDR3 motif enrichment, Vβ gene usage, and CDR3 length (see Methods). Of these, 92 GLIPH2 groups (86%, ☆) were linked to recognition of Mtb-infected macrophages and 15 (14%, ⊡) to MTB300 peptides but not infected macrophages (**Figs. 3A and 3B**). We searched for these TCRβ sequences in the immune epitope database (IEDB, iedb.org) (Chronister et al., 2021) and removed GLIPH2 groups that contained TCRs with ≥ 97% CDR3β homology with TCRs annotated as specific for viral antigens (**Fig. S3A**). Finally, we eliminated 8 GLIPH2 groups that contained TCRs from non-LTBI participant responses to infected macrophages (**Fig. S3A**).

**Figure 3.**
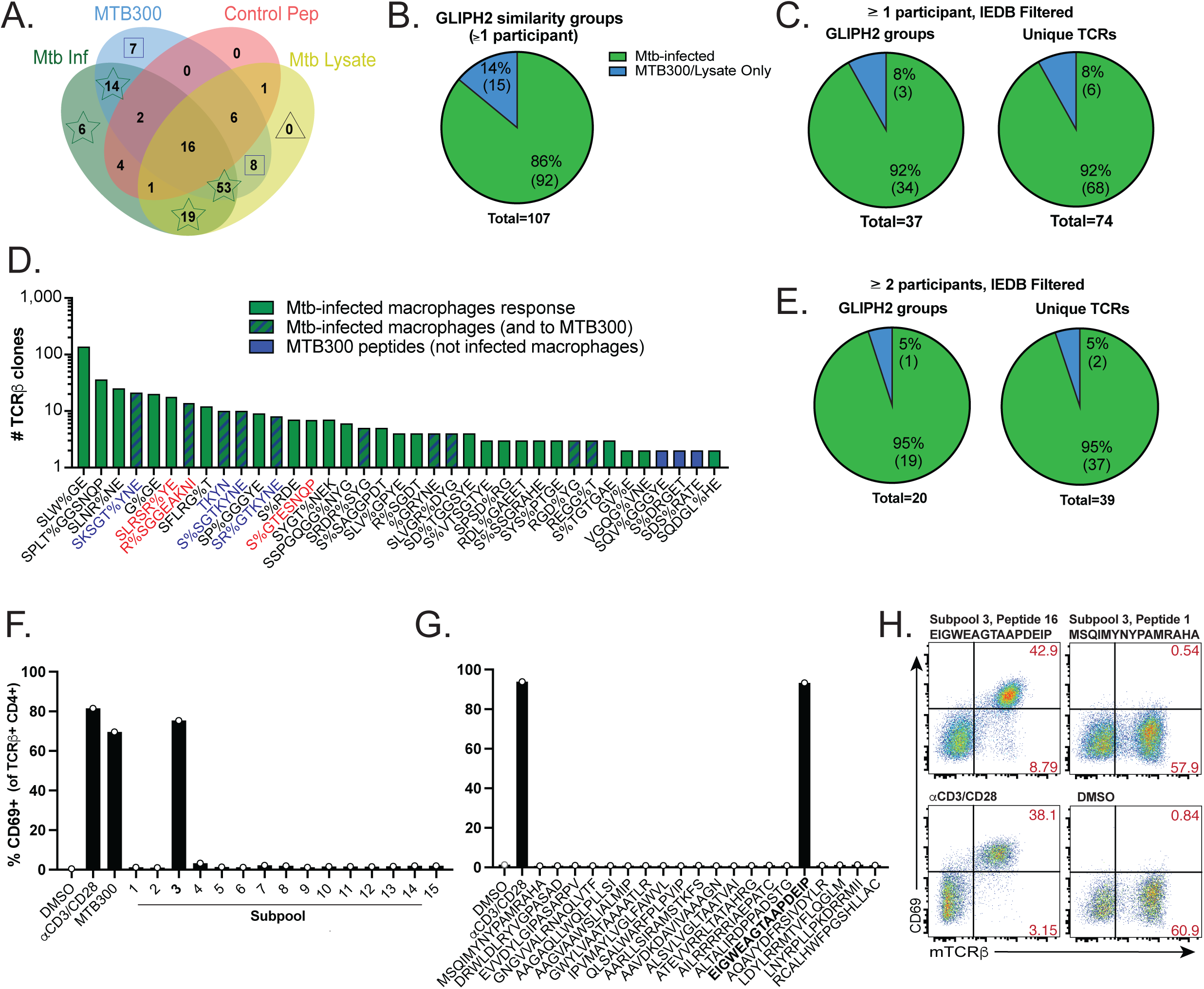
GLIPH2 refines estimates of Mtb-specific CD4^+^ T cell recognition of infected macrophages. **(A)** Venn diagram indicating the number of GLIPH2 groups containing TCRs linked to responses to infected macrophages (green, stars), MTB300 (blue, squares), Control peptide megapools (red), or lysate (yellow, triangle). Data were generated from a combined list of TCRs from all experimental conditions (10 experiments; 10 LTBI and 6 non-LTBI participants). **(B)** Pie charts of percent (and number) of remaining GLIPH2 groups that respond to infected macrophages (green; sum of star groups from A) or MTB300 only (blue; sum of square groups from A). **(C)** Pie charts of GLIPH2 groups (left) and corresponding unique TCRβs (right) after removing GLIPH2 groups containing TCRs linked to viral antigen responses. **(D)** Bar graph of GLIPH2 groups (x-axis) estimated to be Mtb-specific, rank-ordered by sum of highest number of TCR copies per condition (y-axis). Responses to infected macrophages (green), MTB300 only (blue), or both (blue stripes) are indicated. **(E)** Pie charts comparing responses to infected macrophages (green) or MTB300 peptides only (blue) for GLIPH2 groups (left) containing unique TCRβs (right) from at least 2 participants. **(F)** Bar graphs of CD69 expression of ‘TKYN’ TCR-transduced SKW-3 cells by flow cytometry (gated on CD4^+^ TCRβ^+^ Live-Dead^Lo^) 18 h after co-culture with APCs loaded with MTB300 megapool, 15 subpools (20 peptides each), or **(G)** individual peptides from subpool #3 from 2 independent experiments. **(H)** Representative flow plots of CD69 and TCRβ expression gated on total CD4^+^ SKW-3 cells after TCR transduction in response to cognate peptide (top left), irrelevant peptide (top right) and controls. See also Supplemental Figure 3.

Of the remaining 37 GLIPH2 groups and 74 unique TCR clonotypes identified, 92% were linked to recognition of infected macrophages and 8% to MTB300 peptides only (**Figs. 3C and 3D**). Three of these motifs (SLRSR%YE, R%SGEAKNI – ‘GEAK’ motif, and S%GTESNQP – ‘TESN’ motifs) were homologous or identical to those published in 3 recent studies of LTBI in South Africa, containing TCRs specific for the Mtb antigens mIHF (Rv1388), CFP10 (Rv3874), and EspA (Rv3616c), respectively (Musvosvi et al., 2023; Huang et al., 2020; Glanville et al., 2017). Their discovery among LTBI donors increased our confidence that our approach enriched for Mtb-specific TCRs. Finally, GLIPH2 groups containing TCRs found among ≥ 2 participants showed a similar trend where the majority (95%) were linked to recognition of infected macrophages (**Fig. 3E**).

To independently examine the antigen specificity of 4 other dominant GLIPH2 groups composed of TCRs from 3 separate individuals (TKYN, SKSGT%YNE, S%SGTKYNE, and SR%GTKYNE), we cloned a representative TCR expressing the CDR3α and β chains ‘CAAVGSSNTGKLIF’ and ‘CASSRSGTKYNEQFF’ using lentiviral transduction (Dezfulian et al., 2023) into the SKW-3 lymphocytic leukemia cell line which lacks TCR expression (Shima et al., 1986). Epitope screening of SKW-3 cells containing the TKYN TCR in response to EBV-transformed B cells loaded with the MTB300 megapool, or subpools of 20 peptides each, led to CD69 expression only in response to subpool 3 (**Fig. 3F**). Screening the individual peptides from subpool 3 revealed CD69 expression only in response to the ‘EIGWEAGTAAPDEIP’ peptide from EccE3 (Rv0292), a subunit of the Esx3 type VII secretion system (Famelis et al., 2019) (**Figs. 3G and 3H**). These results highlight the enrichment for Mtb-specific TCRs with our approach, and the dominant recognition of Mtb-specific CD4^+^ T cells to infected macrophages.

### TCRs from additional LTBI cohorts enhance the ability of GLIPH2 to distinguish Mtb-specific TCR clonotypes

Our sample size of 16 participants (10 LTBI, 6 non-LTBI) limited our ability to ensure GLIPH2 groups were robust by using HLA associations or containing TCRs from >2 participants (Musvosvi et al., 2023). To enrich our dataset, we added TCRs from > 100 participants in 3 recent publications of TCRs for CD4^+^ T cells activated in response to Mtb antigens in the form of Mtb lysate, MTB300 peptides, or overlapping peptide libraries spanning the ESAT6 and CFP10 antigens (Glanville et al., 2017; Huang et al., 2020; Musvosvi et al., 2023). Using this combined dataset, we focused only on GLIPH2 groups containing at least one unique TCR from LTBI participants in Cleveland, significant Fisher’s exact test, Vβ usage, CDR3β length, and HLA association scores, as described (Musvosvi et al., 2023). We identified 29 GLIPH2 groups with 85 unique TCRs from ≥ 3 participants that did not contain TCRs from control peptide stimulations, non-LTBI participants, or viral antigen-specific TCRs annotated in the IEDB (**Figs. 4A and 4B**). 73% of GLIPH2 groups (and 73% of TCR clonotypes) were linked to recognition of Mtb-infected macrophages **(Figs. 4A and 4B**). 12% of the TCRs within GLIPH groups responded to MTB300 but not infected macrophages, and 15% to lysate but not MTB300 or infected macrophages (**Fig. 4B**). GLIPH2 groups linked to recognition of infected macrophages contained more TCR copies per sample (**Fig. 4C**). In the combined analysis, 8 of these GLIPH2 motifs (SLRSR%YE, R%SGGEAKNI and RA%GGEAKNI – ‘GEAK’ motifs, SPGTESN%P, SLGTESN%P, and S%GTESNQP– ‘TESN’ motifs, SRDN%P and ALFG) were previously published (Musvosvi et al., 2023; Glanville et al., 2017; Huang et al., 2020) revealing specificity for mIHF, CFP10, EspA, and two Mtb antigens yet to be defined, respectively (**Fig. 4C**). Interestingly, the ‘GEAK’, SRDN%P, and ALFG motifs were enriched among “non-progressors” to active TB from the Adolescent Cohort Study in South Africa (Musvosvi et al., 2023).

**Figure 4.**
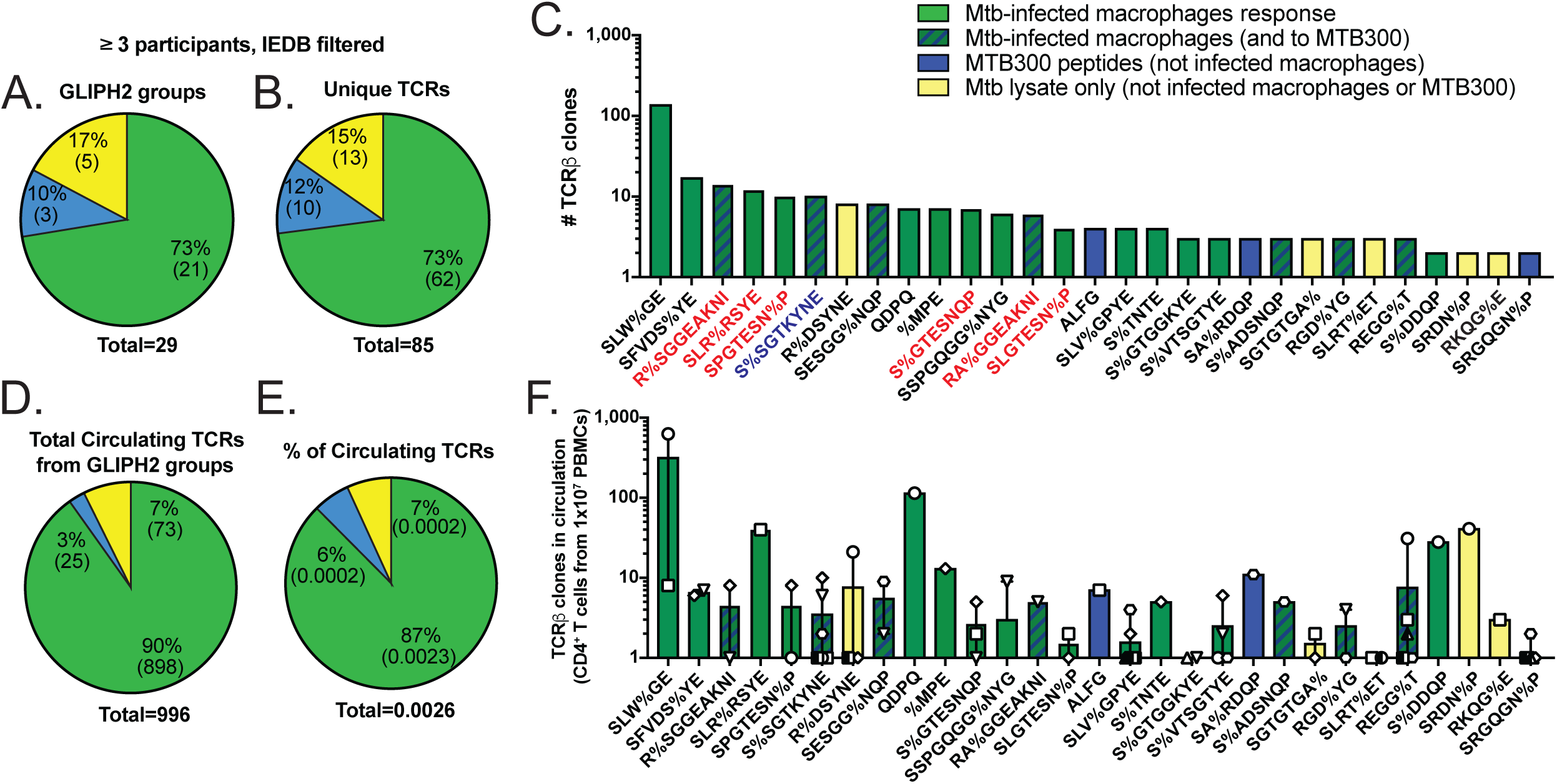
TCRs from additional LTBI cohorts enhance the ability of GLIPH2 to distinguish Mtb-specific TCR clonotypes. **(A)** Pie charts comparing percent (and number) of GLIPH2 groups or **(B)** unique TCRβs (right) linked to a response to Mtb-infected macrophages (green), MTB300 only (blue), or lysate only (yellow) from ≥ 3 participants in the combined TCR dataset. **(C)** Bar graph of GLIPH2 groups estimated to be Mtb-specific (x-axis) from ≥ 3 participants, rank-ordered by sum of highest number of TCR copies per condition (y-axis). Responses to infected macrophages (green), MTB300 only (blue), both (blue stripes), or lysate only (yellow) are indicated. GLIPH2 groups containing TCRs annotated as Mtb antigen-specific in IEDB, or from our peptide screen, are in red and blue font, respectively. **(D)** Pie charts comparing total or **(E)** percent of total circulating TCRβs for CD4^+^ T cells from 10×10^6^ unstimulated PBMCs after cross-referencing CDR3β sequences from Cleveland participants from GLIPH2 groups. Each dot represents TCRβ count from a separate participant. **(F)** Bar graph of mean circulating frequency of Cleveland participants’ TCRβ clonotypes (symbols) for each GLIPH2 group after cross-referencing with unstimulated PBMCs. **(G)** Bar graphs of mean CDR3β (top) and CDR3α (bottom) lengths of TCRs from Cleveland participants within GLIPH2 groups from the combined dataset. **(H)** Sequence logo plots show the probability of each amino acid for CDR3β (top) and CDR3α (bottom) motifs of six GLIPH2 groups common in the initial and combined analyses, created using WebLogo3. The number of CDR3 sequences used for each plot is indicated (top-right). See also Supplemental Figure 3.

To estimate the natural circulating frequency of each TCR from our participants within these GLIPH2 groups, we performed ultra-deep TCRβ sequencing of bulk peripheral blood CD4^+^ T cells after immunomagnetic CD4 selection from 10×10^6^ PBMCs per donor. We then cross-referenced the CDR3β sequences from each of the 29 GLIPH2 groups with those in circulation. Overall, 90% of the total TCRs enumerated in circulation from GLIPH2 groups were linked to recognition of infected macrophages, representing a circulating frequency of 0.0023% (**Figs. 4D and 4E**). Interestingly, some of the GLIPH2 groups containing TCR clonotypes that responded only to MTB300 peptides (ALFG, SA%RDQP) or lysate (R%DSYNE, SRDN%P) had comparable frequencies in circulation to TCRs from GLIPH2 groups linked to recognition of infected macrophages (**Fig. 4F**). All but 6 GLIPH2 groups demonstrated consistent CDR3α sequence homology, indicating a high likelihood they respond to the same peptides. GLIPH2 groups with “inconsistent” CDR3α homology included SLW%GE, REGG%T, RGD%YG, QDPQ, S%TNTE, and S%DDQP. In contrast, 56 GLIPH2 groups were categorized as not specific for Mtb antigens (**Figs. S3B and S3C**). In summary, most GLIPH2 groups enriched for Mtb-specific TCRs contained homologous CDR3α and β sequences and were linked to recognition of infected macrophages.

### Single-cell transcriptomics reveals distinct phenotypic clusters of memory CD4^+^ T cell responses to Mtb-infected macrophages

To explore the effector responses of memory CD4^+^ T cells to Mtb-infected macrophages, we performed scRNAseq 16-18h after *ex vivo* co-culture with infected macrophages (± treatment with lysate) in parallel with scTCRseq (**Fig. 1A**). We did not evaluate the gene expression of MTB300-treated samples to avoid comparing peptide-stimulated T cells with responses to antigens naturally processed and presented by infected or lyste-treated macrophages. An integrated dataset was generated from 157,462 high-quality CD4^+^ T cells using Seurat V5 (Satija et al., 2015) after removing transcripts and cells that did not meet quality control thresholds, and regression of genes that could bias cluster formation (i.e. TCR V and J genes). 19 unsupervised cell clusters were generated and visualized using Uniform Manifold Approximation and Projection (UMAP) (**Fig. S4A**). The three smallest clusters were removed for quality control due expression of macrophage markers CD11c (ITGAX) and MRC1, or low cell numbers (see Methods) (**Figs. S4B-D**), and we analyzed the remaining 16 clusters (**Fig. 5A**).

**Figure 5.**
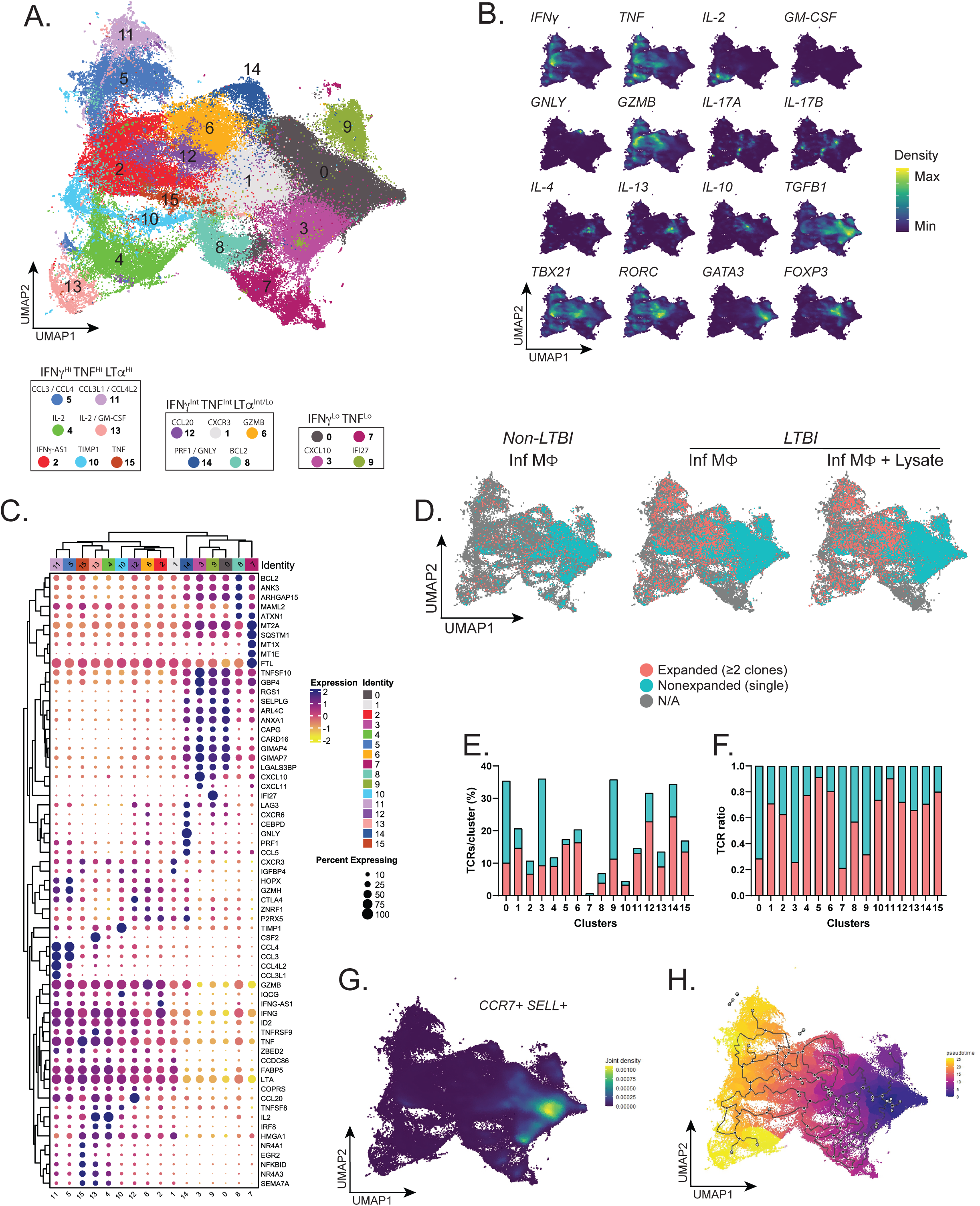
Single-cell transcriptomics reveals distinct phenotypic clusters of memory CD4^+^ T cell responses to Mtb-infected macrophages. **(A)** UMAP visualization plot including Louvain clustering of 157,462 cells, flow-sorted based on expression of CD4 and AIMs, from 7 LTBI participants (12 samples, including memory CD4^+^ T cells in co-culture with Mtb-infected macrophages ± lysate) and 6 non-LTBI participants (6 samples, memory CD4^+^ T cells in co-culture with Mtb-infected macrophages) after integration and QC. **(B)** Kernel density estimation of gene transcripts for T helper subset genes projected onto UMAP plot. Density metrics values were reduced to max/min scale. **(C)** Heatmap with hierarchical clustering showing top 5 DEGs (left) for each cluster (top and bottom), conserved across treatment groups. **(D)** Split UMAP plots for experimental groups showing mapping of all TCRs. Expanded (≥2 copies) and non-expanded (single) TCR clonotypes are shown in red and blue, respectively. **(E)** Representative stacked bar plots showing percent clonally expanded versus non-expanded TCRs in LTBI participant samples, normalized to each cluster’s total cell number and **(F)** total number of TCRs. **(G)** UMAP plot showing joint density estimation for plot for CCR7 and SELL transcripts in the integrated dataset. **(H)** UMAP plot illustrating the single-cell trajectory and pseudotime analysis of CD4+ populations within the integrated dataset. Cell fates (gray circles), transition states (black circles), proximity to (purple), and remoteness from (yellow) the root are indicated. See also Supplemental Figure 4.

To define the phenotypic composition of each T cell cluster, we used the combination of gene density visualization for canonical CD4^+^ T cell markers and the top 5 conserved differentially expressed genes (DEGs) for each cluster (**Figs. 5B and 5C and Fig. S4E**). Higher expression of *IFNG* and *TNF*, canonical Th1 cytokines, were observed in 13 of 16 clusters, with maximum levels of expression in clusters 2, 4, 5, 10, 11, 13, and 15 (**Fig. 5C and Fig. S4E**). High expression of *CXCR3*, together with *ID2*, *LTA* (lymphotoxin-α), and *GZMB* (granzyme B) were also observed in the same 13 clusters (**Fig. 5C and Fig. S4E**). In addition to *IFNG* and *TNF*, *IL-2* expression was prominent in clusters 4 and 13, with cluster 13 also showing strong and unique expression of *CSF2* (GM-CSF). Chemokines *CCL3* and *CCL4*, which are associated with immune cell recruitment (Castellino et al., 2006), were preferentially expressed in clusters 5 and 11, while *CCL3L1* and *CCL4L2* were co-expressed in cluster 11. In contrast, cells in clusters 1, 6, 8, 12, and 14 showed intermediate or low expression of *IFNG*, *TNF*, and *IL2*. In addition to expressing *GZMB*, the top DEGs for cells in cluster 14 were the cytotoxic molecules *PRF1* (perforin) and *GNLY* (granulysin), indicating cytotoxic CD4⁺ T cell responses. Although cluster 6 expressed the highest levels of *GZMB*, clusters 12 and 8, were enriched for *CCL20* and *BCL2*, respectively. Cluster 1 was enriched for expression of *CXCR3*, but also contained expression of several other cytokines, including *IL10*, *IL17A*, and *IL17F* (**Figs. 5A-C**). Clusters 0, 3, 7, and 9 expressed little to no *IFNG* or *TNF*, and instead expressed the type I and II IFN response genes *IFI27* and *CXCL10*, respectively, along with *TGFB1* and *IL10* (**Fig. 5B**). Together, these data indicate a distribution in the phenotypes of CD4^+^ T cell responses to Mtb-infected macrophages ranging from canonical Th1 functions, cytotoxicity, GM-CSF, and chemokine secretion to the expression of regulatory cytokines and response to interferons.

### CDR3 mapping distinguishes the effector functions of Mtb-specific TCR clonotypes that recognize infected macrophages

To examine the functions of clonally expanded CD4⁺ T cells in response to infected macrophages (± treatment with lysate), we first visualized the mapping of expanded TCRs (≥2 copies) compared to non-expanded TCRs (1 copy) from LTBI and non-LTBI participants (**Fig. 5D**). Among all groups, single TCR clonotypes were densely localized on the right side of the UMAP plot, within clusters 0, 3, 7, and 9 (**Figs. 5D-F**), which lacked expression of *IFNG*, *TNF*, and *LTA* (**Figs. 5A-C**). The transcriptional profiles in clusters 0 and 3, including co-expression of *CCR7* and *SELL* (CD62L), suggested minimal differentiation and a central memory T cell phenotype (**Fig. 5G**). Pseudotime projection anchored within clusters 0 and 3 demonstrates that clusters 4 and 13, contained the most differentiated CD4^+^ T cells, followed by clusters 11 and 5 (**Figs. 5G and 5H**), coinciding with the expanded TCRs (**Figs. 5D and 5E**). This distribution pattern highlights the importance of canonical CD4^+^ T cell effector molecules IFNγ, TNF, LTα, IL-2, and GM-CSF, cytotoxic molecules granulysin, perforin, and granzyme B, and chemokines CCL20, CCL3, CCL4, CCL3L1, and CCL4L2 in response to infected macrophages.

To examine the relationship between CD4⁺ T cell phenotypes and their capacity to recognize infected macrophages, we mapped TCRs from relevant GLIPH2 groups onto single-cell transcriptomics data. By identifying TCRs with CDR3α sequence homology (differing by ≤ 2 amino acids) that formed motifs, we categorized GLIPH2 groups as "consistent", if displaying high CDR3α and CDR3β homology, and "inconsistent" when displaying greater CDR3α variation (**Fig. S5A**). We focused on TCRs from consistent GLIPH2 groups (**Fig. 4H**). We mapped previously annotated Mtb-specific CDR3β motifs, TCRs likely to be Mtb-specific, and their corresponding GLIPH2 groups to the UMAP plot (**Figs. 6A-6D**). GLIPH2 groups with inconsistent CDR3α homology mapped with a different distribution pattern, enriched in clusters 0, 1, 3, 5, 6, and 12 (**Figs. S5A and S5B**). Although fewer clonotypes responded only to MTB300 and/or lysate, they distinctly mapped to clusters 0, 3, and 14 (**Figs. 6E and 6F**).

**Figure 6.**
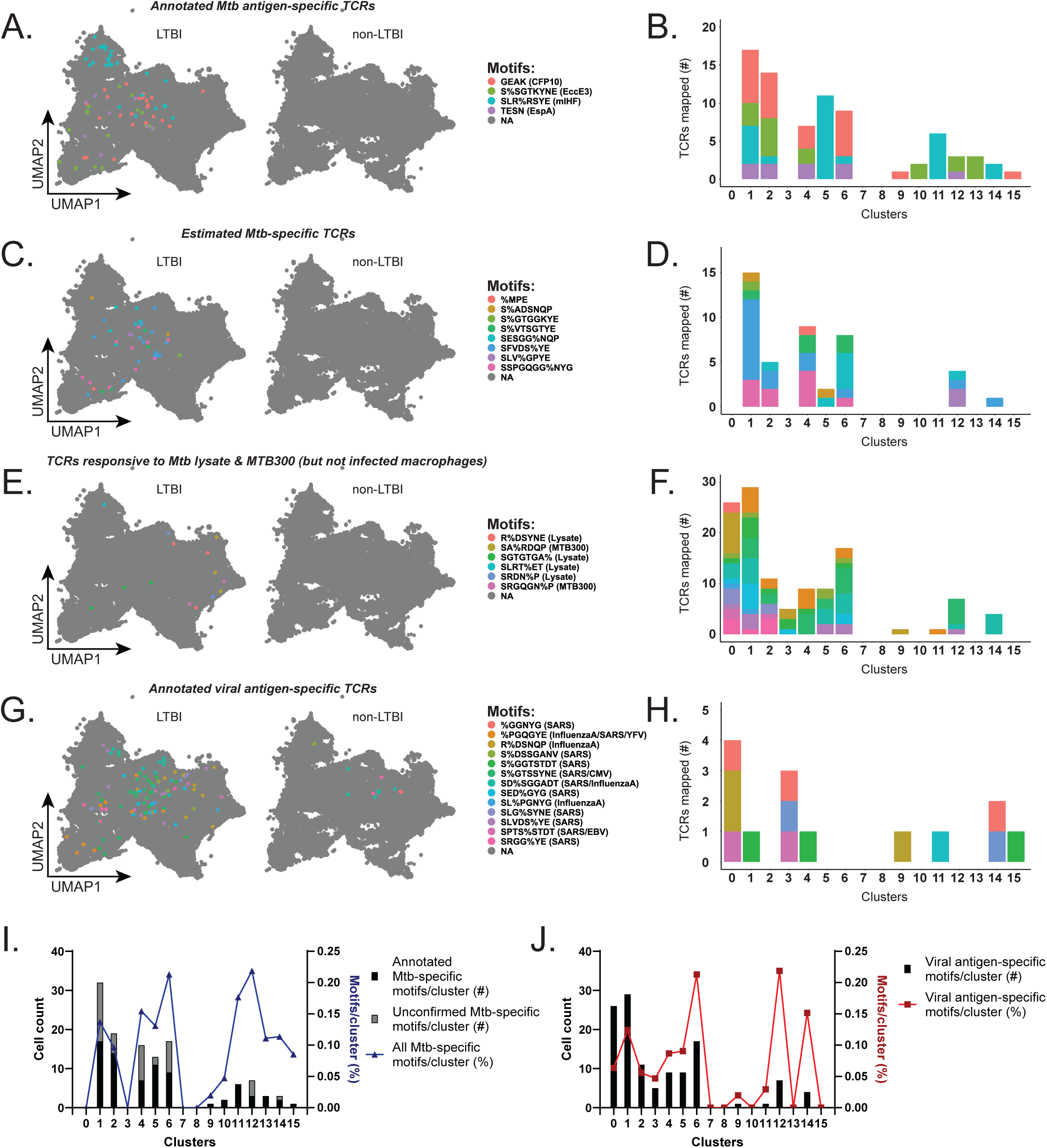
CDR3 mapping distinguishes effector functions of Mtb-specific TCR clonotypes that recognize infected macrophages. **(A)** UMAP plot with split view (based on LTBI status) with mapping of TCRs from listed GLIPH2 global motifs and **(B)** stacked bar plots showing their numbers per cluster for previously annotated Mtb antigen-specific TCRs. **(C)** UMAP plots with mapping of TCRs estimated to be Mtb-specific from listed GLIPH2 groups and **(D)** stacked bar plots showing their numbers per cluster. **(E)** UMAP plots with mapping of TCRs responsive to MTB300 or lysate stimulation, but not infected macrophages, from listed GLIPH2 groups and (F) stacked bar plots showing their numbers per cluster. **(G)** UMAP plots with mapping of annotated viral antigen-specific TCRs from listed GLIPH2 groups and **(H)** stacked bar plots showing their numbers per cluster. **(I)** Total copy numbers (left axis) and percentage (right axis) of cells per cluster mapping TCRs ([clone count / total cells per cluster] x 100) from known or estimated Mtb-specific GLIPH2 groups that recognized infected macrophages and **(J)** for annotated viral antigen-specific TCRs. See also Supplemental Figure 5.

To further delineate the phenotype of antigen-specific CD4^+^ T cell responses to infected macrophages, we evaluated GLIPH2 groups containing at least one TCR with 100% CDR3β homology to viral antigen-specific TCRs annotated in the immune epitope database. Clusters 0, 1, 6, 12, and 14 were enriched for viral antigen-specific TCRs in both LTBI and non-LTBI (**Figs. 6G and 6H**). TCRs from GLIPH2 groups with inconsistent CDR3α homology mapped to the same clusters as viral antigen-specific responses (**Figs. S5C and S5D**). TCR mapping was also represented as the percentage of total cells per cluster to reduce TCR assignment bias due to differences in cell numbers per cluster (**Figs. 6I and 6J and Figs. S5E and S5F**). Enrichment of Mtb-specific TCRs was found among clusters 2, 4, 5, 10, 11, 13, and 15 whereas a higher proportion of viral antigen-specific TCRs mapped to clusters 0, 3, 6, 12, and 14 (**Figs. 6I and 6J, and Fig. 7A**). Interestingly, Mtb-specific TCRs were enriched among clusters 11 and 13, the two most differentiated CD4^+^ T cell clusters in response to infected macrophages (**Fig. 5H**). These data highlight that in response to infected macrophages, Mtb-specific TCR clonotypes express distinct effector functions.

**Figure 7.**
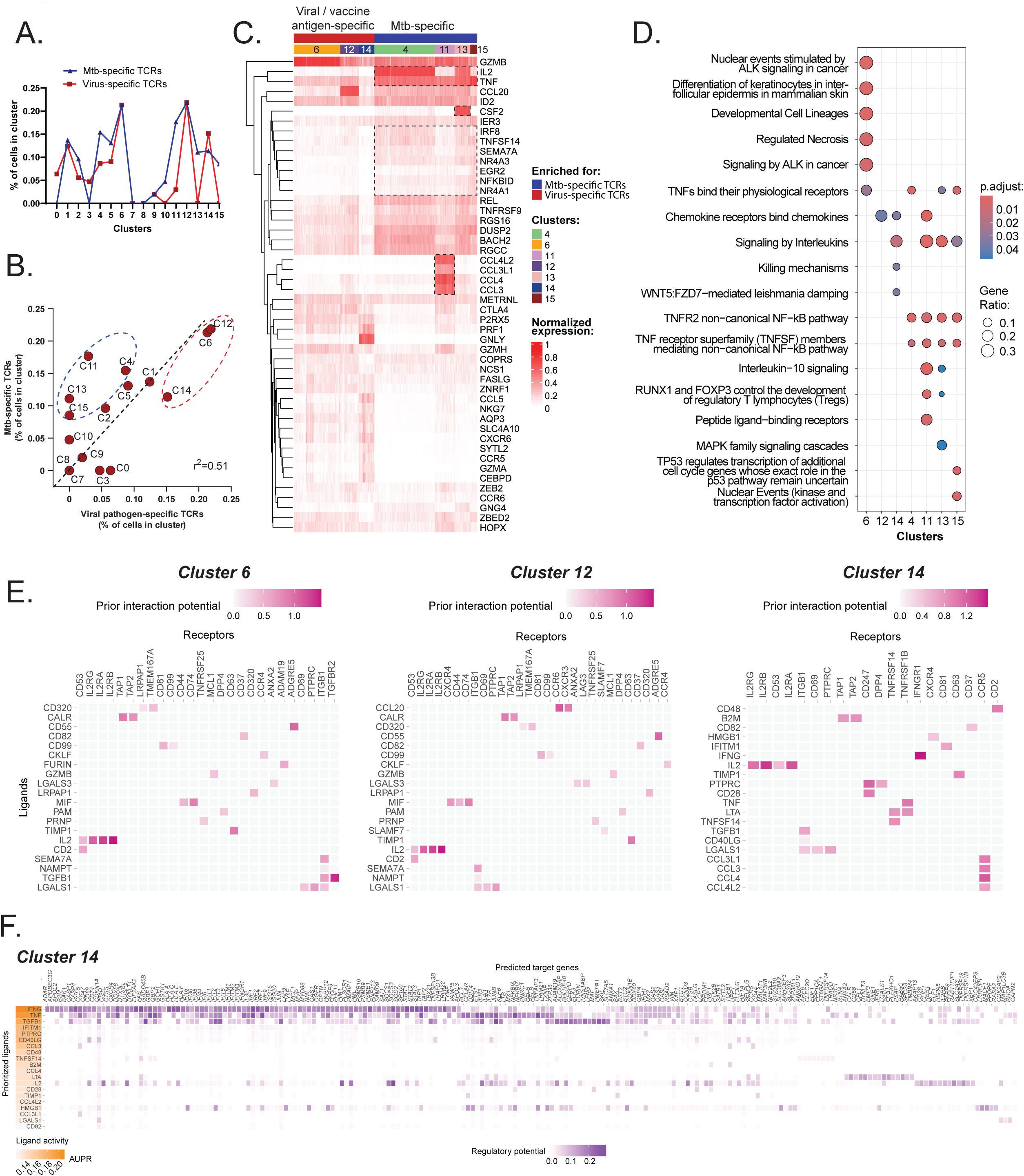
Mtb-specific CD4^+^ T cells feature two signature gene sets in response to infected macrophages. **(A)** Overlaid line graph and **(B)** correlation plot of percent T cells per cluster that mapped TCRs estimated to be Mtb-specific (blue) and viral antigen-specific (red). Linear regression (black dashed line) correlation was estimated using Pearson correlation coefficient squared (r^2^). Red and blue dashed circles identify UMAP clusters with highest enrichment of TCRs estimated to be Mtb or viral antigen-specific, respectively. **(C)** Heatmap showing top 10 DEGs normalized to the individual maximum expression for clusters 4, 6, 12, 14, 11,13, and 15. Gene expression patterns common to cluster enriched for Mtb-specific TCRs (4,11, 13, and 15) are outlined. **(D)** Dot plot showing Reactome pathway overrepresentation analysis using the lists of genes with Log_2_FC>1 for UMAP clusters 4, 6, and 11-15. The list of top 7 most significant pathways is arranged based on the GeneRatio (number of input genes associated with a Reactome term / total number of input genes). **(E)** Summary of receptor-ligand pairs identified by NicheNet analysis, estimating the cell-cell communication between “sender” clusters 4, 11, 13, or 15 (combined) and “receiver” cluster 6 (left), cluster 12 (middle), and cluster 14 (right). **(F)** Heatmap showing downstream signaling genes estimated to be linked to cell-cell communication between the cluster 4, 11, 13, or 15 ligands (combined) and cells in cluster 14. The area under the precision-recall curve (AUPRC) was used to rank the ligand activity of senders on responders (left). See also Supplemental Figure 6.

### Mtb-specific CD4^+^ T cells feature two signature gene sets in response to infected macrophages

To determine the signature gene expression profiles linked to Mtb-specific recognition of infected macrophages, we examined the correlation between the frequencies of Mtb and viral antigen-specific TCRs from corresponding GLIPH2 groups with consistent CDR3α and β homology that mapped to each cluster. Normalized TCR mapping analysis revealed that clusters 4, 11, 13, and 15 were enriched with both established and probably Mtb-specific TCRs and GLIPH2 groups (**Figs. 7A and 7B**). Clusters 6 and 12 mapped the greatest frequencies of both Mtb and viral antigen-specific TCRs as a proportion of total cells, and cluster 14 contained the highest frequency of viral antigen-specific TCRs (**Fig. 7B**). While cluster 11 featured some viral antigen-specific TCRs, it contained a 6-fold greater proportion of Mtb-specific clonotypes. Mtb-specific TCRs were also over-represented in cluster 4, which expressed a highly differentiated profile (**Fig. 5H**) and its phenotype closely resembled that of cluster 13 (**Figs. 5C**). The same distributions were observed if TCRs from all 29 GLIPH2 groups were included for analysis (**Fig. 4C and Figs. S5G and S5H**). Therefore, we focused on clusters 4, 6, and 11-15 for differential gene expression analyses.

Clusters enriched with Mtb-specific TCRs included preferential expression of *CCL3*, *CCL4*, *CCL3L1*, and *CCL4L2* (cluster 11), *IL-2* (clusters 4 and 13), *GM-CSF* (cluster 13), and *IRF8, TNFSF14, SEMA7A, NR4A3, NR4A1,* and *NFKBID* (clusters 4, 13, and 15) as top DEGs (**Fig. 7C**). To examine the pathways involved, we performed Reactome pathway over-representation analysis using the top DEGs (Log_2_FC > 1) for clusters 4, 6, and 11-15. Despite containing distinct gene expression signatures, all clusters enriched for Mtb-specific TCRs (i.e., 4, 11, 13, and 15) shared a biological theme centered around non-canonical NFκB pathway signaling mediated by TNF superfamily members or TNFR2 signaling (**Figs. 7D and Fig. S6A**). Clusters with the highest ratio of Mtb to viral antigen-specific TCR mapping, (i.e. clusters 11 and 13), were found to activate T_Reg_ development and IL-10 signaling pathways. The latter is of particular interest, since we and others have previously shown that human and murine MDMs produce IL-10 upon infection with Mtb (Gail et al., 2023; Moreira-Teixeira et al., 2017). Pathways for clusters 6 and 14 were dominated by cytotoxic responses (**Fig. 7D and Fig. S6B**). Together, these data identify pathways common and unique to clusters enriched for Mtb-specific TCR clonotypes.

Finally, to test the hypothesis that Mtb-specific clonotypes express factors that contribute to the recruitment and activation of other memory T cells to control infection, receptor-ligand pairs were evaluated by analysis with NicheNet (Browaeys et al., 2020). Using the DEGs from T cells enriched with Mtb-specific TCRs (clusters 4, 11, 13, and 15), as potential “sender” ligands, we identified IL-2 signaling through CD53, and IL-2Rα,β, or γ, as the ligand-receptor pair common to clusters 6, 12, and 14 with a dominant downstream signaling pattern in all 3 clusters, while GM-CSF was not linked to any receptor-ligand interaction (**Figs. 7E and Fig. S6C**). IL-2 signaling correlated with expression of *IL2RA*, *IL2RB*, and *BCL2L1*, *MYC*, *PIM1*, *SOCS3*, *DUSP5*, *GZMB*, and *PRF1*, indicating a cytotoxic effector program, and *BHLHE40*, a strong regulator of IL-10 transcription (Huynh et al., 2018) (**Fig. 7F and Fig. S6C**). CCL3, CCL4, CCL3L1, and CCL4L2 signaling through CCR5, IFNγ signaling through IFNγR1, TGFβ signaling through ITGB1, and TNF and LTα signaling through TNFR2 also represented candidate ligand-receptor interactions with cluster 14 (**Figs. 7E and 7F**). Remarkably, each of these cytokine and chemokine ligands represented top DEGs preferentially expressed by the clusters enriched with Mtb-specific TCRs, except for TGFβ, which is expressed by T cells in clusters 0 and 3 (**Figs. 5A-C and Fig. 7C**). TFGβ was also shown to be expressed by Mtb-infected MDMs (Gail et al., 2023). These data indicate that the top cytokines and chemokines expressed by Mtb-specific CD4^+^ T cell clusters have the potential to provide a narrow set of recruitment and activation signals to other memory T cells and could serve a protective role against Mtb infection.

## Discussion

In this study, we present the first analysis describing human memory CD4^+^ T cell responses to autologous macrophages infected with virulent Mtb using single-cell TCR sequencing and transcriptomics. The majority of memory CD4^+^ T cells from individuals with LTBI recognized infected macrophages, but a subset responded only when exogenous antigens were added. Among 10 individuals with LTBI, we find 73-90% of the Mtb-specific TCR clonotypes were linked to recognition of infected macrophages. We discovered several of these TCRs from multiple individuals to contain specificity for Mtb antigens. We focused on clonally expanded TCRs (≥ 2 copies) and TCRs containing CDR3 motifs overrepresented among multiple individuals. We also used GLIPH2 to segregate TCRs specific for other pathogens, indicated by a response to control peptide stimulations or those annotated as specific for viral antigens in the IEDB. While this approach could unintentionally eliminate cross-reactive TCRs that are in fact Mtb-specific, cross-reactivity between Mtb and viral antigens is expected to be rare. Furthermore, the benefit of enriching for Mtb-specific TCRs outweighed the risk of losing a small number of potentially cross-reactive TCRs. The identification of TCRs specific for Mtb antigens from multiple individuals with LTBI, including EccE3 from our screen and CFP10, EspA, and mIHF antigen-specific TCRs annotated in 3 recent studies (Musvosvi et al., 2023; Huang et al., 2020; Glanville et al., 2017) indicates our approach enriched for Mtb-specific CD4^+^ T cells.

T cells sharing similar CDR3 motifs often exhibit overlapping antigen specificity, transcriptional profiles, and cell fate, particularly in the context of infectious diseases (Chen et al., 2023; Wang and Ji, 2023). By mapping TCRs to their gene expression profiles, we discovered Mtb-specific responses linked to recognition of infected macrophages preferentially co-express *IFNG*, *TNF*, *IRF8, TNFSF14, SEMA7A, NR4A3, NR4A1, NFKBID,* and *CCL3*, *CCL4*, *CCL3L1* and *CCL4L2* (**Fig. 7C, cluster 11**), or *IL2* and *CSF2* (GM-CSF) (**Fig. 7C, cluster 13**). Cluster 15 (highest *TNF* expression), cluster 5 (expressing *CCL3* and *CCL4*), and cluster 4 (expressing *IL2*) were also enriched for Mtb-specific TCRs, were in fact adjacent to clusters 11 and 13, and were phenotypically similar but less differentiated based on trajectory analysis (**Fig. 5H**). Clusters 11 and 13 contained the highest enrichment for Mtb-specific TCRs and were the most differentiated T cell clusters in our dataset (**Fig. 5H**). The highest frequencies of all clonally expanded TCRs, and those annotated as viral antigen-specific, mapped to clusters 6, 12, and 14, suggesting bystander T cell activation. These clusters were enriched for cytotoxic gene expression, including *GZMB*, *PRF1*, *GNLY*, and *CCL20*, respectively. The non-expanded TCRs were enriched among clusters 0, 3, 7, and 9, which almost uniformly expressed central memory-like phenotype and regulatory functions, indicating a second program of bystander T cell activation. Although a much smaller subset, TCR clonotypes that responded only to MTB300 and/or lysate (but not infected macrophages) also mapped to clusters 0 and 3. Given their enrichment with Mtb-specific TCRs, and their distinct and differentiated transcriptomics profiles, we conclude the effector phenotypes of clusters 4, 11, 13, and 15 represent Mtb-specific memory CD4^+^ T cell recognition of infected macrophages.

We initially sought to compare Mtb-specific CD4^+^ T cell responses that did or did not recognize infected macrophages but were struck by the large numbers of viral antigen-specific TCR clonotypes activated in response to infected macrophages. We considered both cross-reactivity due to molecular mimicry and non-specific bystander activation of memory T cells. Since humans contain only ∼10^12^ total and ∼10^8^ unique TCR clonotypes available to recognize the >10^15^ foreign peptides (Arstila et al., 1999; Mason, 1998; Sewell, 2012), bystander activation and cross-reactivity of immunological memory extend protection of antigen-specific T cells across multiple infectious diseases (Su et al., 2013; Kundu et al., 2022; Bartolo et al., 2022). Shared epitopes between viral pathogens and Mtb have not been identified. However, one recent study speculated that CD4^+^ T cell clones specific for SARS-CoV-2 antigens in uninfected individuals might gain a memory phenotype from cross-reactivity with commensal microbiota (Bartolo et al., 2022). Exposure to environmental non-tuberculous mycobacteria (NTMs) by participants in our study is likely to generate T cells cross-reactive to homologous NTM proteins (Prasad et al., 2013; Shah et al., 2019). We would expect these responses to express phenotypes similar to antigen-specific T cells elicited by Mtb infection. Yet, we observed highly expanded TCR clonotypes, many of which were found in multiple participants’ samples, that either responded to control peptide stimulation or were already annotated in the IEDB as specific for SARS-CoV-2, influenza A, CMV, or EBV antigens. Although not definitive, these features suggest bystander activation rather than cross-reactivity.

Bystander activation of CD8^+^ T cells during acute viral infection is well-described (Ehl et al., 1997; Tough et al., 1996; Doisne et al., 2004). Although understudied, bystander activation of memory CD4^+^ T cells after vaccination has also been described (Aalst et al., 2017; Causi et al., 2015; Genova et al., 2006). Although reportedly less effective, T cell activation that does not involve TCR-pMHC interactions can occur via cytokines, including IL-2 and IL-15 signaling (Bangs et al., 2006), IL-12 and IL-18 signaling (Srinivasan et al., 2007), or via costimulatory receptors such as 4-1BB (Reithofer et al., 2021) and NKG2D ligation (Balint et al., 2024), or combinations thereof (Aalst et al., 2017). Our results extend earlier observations to TB, where we find antimicrobial functions among clonally expanded bystander memory CD4^+^ T cells but regulatory functions (e.g., IL-10, TGFβ production) among non-expanded bystander TCR clonotypes. In addition to their antimicrobial functions, our data suggest Mtb-specific CD4^+^ T cells participate in the recruitment and activation of other memory T cells which may underpin the expression of two or more effector programs by Mtb-specific T cells. The cell-cell interaction analysis suggests antigen-specific CD4^+^ T cells interact with bystander-activated CD4^+^ T cells through IL-2, TNF, IFNγ, and chemokines (CCL3, CCL4, CCL3L1, CCL4L2). Therefore, we infer that bystander T cell activation requires interactions with antigen-specific CD4^+^ T cells as well as infected macrophages, suggesting that antimicrobial immune responses could be amplified from small numbers of Mtb-specific T cells. Finally, the observed activation of non-expanded TCR clonotypes likely represents a regulatory component of the bystander response which may limit inflammation by providing negative feedback.

The enormous diversity of HLA alleles and TCRs in humans is a major barrier to identifying pathogen-specific T cell responses. Diversity in the TCR repertoire is driven by variation in HLA alleles between and within individuals, and by differences in the repertoires of Mtb peptides presented to T cells upon infection. As a result, clonal populations of T cells that contain Mtb-specific TCRs are typically unique to a single individual. TCR grouping algorithms like GLIPH2 (Huang et al., 2020) identify clonotypes that contain homologous CDR3β motifs present in samples from multiple individuals who share a common HLA allele and were shown to have specificity for the same peptide epitope. Using lists of Mtb-reactive TCRs from both Cleveland and South African participants, including from the Adolescent Cohort Study (Musvosvi et al., 2023), we identified four GLIPH2 groups which were identical to those from the “non-progressors” who did not progress to active TB (R%SSGEAKNI, S%GTESNQP, ALFG, and SRDN%P). Another four GLIPH2 groups were identical or homologous (differing by 1-2 amino acids) to South African samples in two other studies (Glanville et al., 2017; Huang et al., 2020), including SLR%RSYE, SPGTESN%P, RA%GGEAKNI, SLGTESN%P). All GLIPH2 groups in common with the Adolescent Cohort Study were from non-progressors, suggesting our participants with stable LTBI contained protective TCR clonotypes.

Although a majority of TCRs confirmed or estimated to be specific for Mtb antigens were linked to recognition of infected macrophages, at least 10% responded only when MTB300 peptides or lysate were added. Recent data from the mouse model of TB similarly demonstrated not all Mtb-specific T cells generated during TB are able to recognize infected macrophages (Yang et al., 2018; Patankar et al., 2020). Such Mtb-specific T cells could be specific for peptides that are presented by infected macrophages but might interact with suboptimal binding parameters (e.g. TCR dissociation rate or avidity) and thus inefficiently recognize infected macrophages. It is also possible infected macrophages do not present the cognate peptides for a subset of TCRs. Bacterial regulation of antigen expression (Bold et al., 2011; Moguche et al., 2017), the export of Mtb proteins (Srivastava et al., 2016), or the timing of presentation of cognate Mtb antigens by MDMs in our ex vivo system are all possibilities. Based on recently published work, differences in antigen processing and presentation between DCs in MLNs and macrophages in the lung could account for the priming of T cells that do not recognize infected macrophages (Ewanchuk and Yates, 2018). Upon infection, DCs in MLNs likely present a broader repertoire of Mtb peptides than infected macrophages and therefore prime a broad repertoire of antigen-specific T cells. Supporting this hypothesis, differences in phagocyte oxidase (NOX2) activity was shown to directly affect thiol transferases and cysteine cathepsin activity, and therefore the repertoire of peptides presented in the context of MHC-II (Allan et al., 2014; Rybicka et al., 2010). The regulation of these cathepsins by NOX2 is likely more important to epitopic preservation in lung macrophages where the consequences of activating T cells and promoting inflammation can be severe, affecting gas exchange. It is also logical that to fight infection, the immune system would evolve ways to recruit and activate other memory T cells activation that depend on (but don’t entirely comprise) antigen-specific responses, while preventing the activation of all surround T cells in tissue. Individuals with recent Mtb infection and IGRA conversion represent a state of higher risk for active TB, while exposed individuals who remain IGRA-negative, or have longstanding stable IGRA-positivity are associated with greater protection from active TB (Andrews et al., 2012; Kim et al., 2024). Preferential activation and expansion of Mtb-specific T cells that recognize infected macrophages in stable LTBI may explain the responses in our study. We posit that individuals who possess a greater risk of active TB have a higher proportion of memory CD4^+^ T cells that do not efficiently recognize infected macrophages.

### Limitations of the study

Our study is the first to examine primary human memory CD4^+^ T cell recognition of Mtb-infected macrophages using single-cell transcriptomics yet has limitations. Technical limitations prevented us from sequencing TCRs from every T cell. However, the combined use of GLIPH2, and peptide stimulations allowed us to focus on common CDR3 motifs to estimate the Mtb-specific response, separating it from bystander responses. We recognize drawbacks to focusing exclusively on TCRs grouped by GLIPH2, including the possibility of missing “ungrouped” Mtb-specific clonotypes. To address this, we leveraged the published South African datasets to improve GLIPH2 grouping by increasing the numbers of participant and TCRs. Our initial analysis of clonally expanded TCRs contained many that were viral antigen-specific. Like infection, Mtb lysate has adjuvant properties and could have overestimate the lack of recognition of infected macrophages when used as a comparator. Our *ex vivo* assay uses virulent Mtb and a standard macrophage infection model. While the timing of antigen expression, processing, and presentation could be different *in vivo,* most TCR clonotypes that responded to Mtb peptides or lysate also recognized infected macrophages, including TCRs previously annotated as Mtb-specific. Another limitation was the number of participants included in scRNAseq analyses. However, the trade-off was the ability to assess multiple conditions using costly high throughput scRNAseq. The specificities of many TCRs from our study are still yet to be determined. However, this study provides a link between these TCRs and recognition of infected macrophages (or lack thereof), enabling future screens for antigen specificity as technology makes the process more efficient.

## Materials and Methods

### Study Participants

10 healthy participants with LTBI and a median age of 35 years (range 23-68; 6 male, 4 female), representing African, Asian, Caucasian, and Hispanic ethnicities volunteered based on self-identified history of latent Mtb infection (LTBI). LTBI status was determined based on positive results of either a tuberculin skin test (TST) of at least 10mm or an IFNγ release assay (IGRA) or both. No LTBI participants had a history of active TB disease or symptoms suggestive of current disease. Eight of 10 participants previously spent time in TB endemic areas, and 5 participants previously received isoniazid or rifampin antibiotic prophylaxis at least 3 years prior to participating. Five individuals had remote PPD conversion and had never received antibiotic prophylaxis. Five individuals received the BCG vaccine as infants (>30 years prior to participating). Seven healthy volunteers without LTBI with a median age of 29 years (range 23-53; 4 male, 3 female) served as controls, including anonymized leukapheresesis products from 5 participants purchased from AllCells (Alameda, CA, USA). The negative LTBI status of participants was verified through QuantiFERON-TB Gold Plus (Qiagen, Hilden, Germany). Informed consent was obtained for all participants. The study included both male and female participants and the findings are expected to be relevant to both sexes. No correlation between participant sex and T cell activation was observed, although sample sizes were limited. All protocols involving human subjects were approved through the Institutional Review Board of University Hospitals Cleveland Medical Center. Informed consent was obtained for all participants. Protocols involving lentiviral transduction of human TCR genes were approved by the Institutional Biosafety Committee of Case Western Reserve University.

### Generation of monocyte-derived macrophages

Following blood draw, Ficoll-Paque PLUS (GE Healthcare, Uppsala, Sweden) was used to underlay diluted whole blood using a Pasteur pipet. Following centrifugation, the peripheral blood mononuclear cell (PBMC) layer was aspirated using a transfer pipet. The PBMCs were then washed with Ca^++^ and Mg^++^ free and pyrogen-free sterile phosphate buffered saline (hereafter termed PBS; Corning; Glendale, AZ, USA) and counted using a hemocytometer after staining with trypan blue (Life Technologies, Grand Island, NY) to exclude dead cells. Once counted, CD14^+^ monocytes were separated from the rest of the PBMCs by positive immunomagnetic selection using anti-human CD14 microbeads (Miltenyi Biotec; Bergisch Gladbach, Germany), per manufacturer instructions. After CD14 selection, both the positive and negative fractions were counted and separately cryopreserved in freezing media (10% DMSO, 90% FBS) in liquid nitrogen.

CD14^+^ monocytes were thawed and rested overnight in complete RPMI1640 media (cRPMI) containing 10% FBS, 2mM of L-Glutamine, Na-Pyruvate, Non-essential amino acids, 0.01M Hepes buffer, 2-mercaptoethanol, and 0.005M NaOH (Gibco, Waltham, MA, USA), as described (Gail et al., 2024). The next morning, CD14^+^ cells were plated in either a 96-well plate (50K cells/well) or a 24 well plate (250K cells/well) and differentiated to macrophages using GM-CSF (25ng/mL) (PeproTech, East Windsor, NJ). Half of media in each well was changed at three days with GM-CSF-containing cRPMI. After a total of six days the macrophages were ready for Mtb infection and media from the entire well was exchanged with cRPMI prior to Mtb infection, as described previously (Gail et al., 2024, 2023).

### Bacterial Culture and Infection

A 1mL aliquot of *Mycobacterium tuberculosis* strain H37Rv (NR-13648, BEI Resources) was thawed and diluted in 9mL of media containing Difco Middlebrook 7H9 broth (BD Diagnostics) supplemented with 10% OADC, 0.2% glycerol, and 0.05% Tween-80, and culture was expanded for use at mid-log phase over 6 days, as described (Gail et al., 2023, 2024). H37Rv was selected since it is a virulent reference strain of Mtb, since lysate was also prepared from H37Rv, and for comparability with the majority of published studies of human macrophage *in vitro* infection models. Cultures were washed and filtered through a 5μm Millex syringe-driven filter unit (Millipore; Duluth, GA, USA) to generate a single-cell suspension. The bacterial count was estimated using the OD_600_ of filtered bacteria in cRPMI, and added to macrophage wells to target a MOI of 4-5, as described (Gail et al., 2024, 2023). After 4 h, wells were washed of extracellular bacteria and incubated in fresh cRPMI overnight. Actual MOI were enumerated from 3-4 separate wells of Mtb-infected macrophages as described (Gail et al., 2024, 2023; Carpenter et al., 2017).

### CD4 memory T-cell co-culture assay

CD14-PBMCs were thawed and rested in cRPMI overnight. Immunomagnetic selection was performed per manufacturer instructions using the Human Memory CD4 T cell isolation kit (Miltenyi Biotec) for negative selection of memory (CD45RA^Lo^) CD4^+^ T cells. AutoMACS Rinsing solution with 5% MACS BSA Stock Solution (Miltenyi Biotec) was used to wash cells, hereafter termed “Rinse Buffer”. After selection, CD4^+^ T cells were then added at ∼4:1 ratio to the infected macrophages in 24 well plates. Ultra-LEAF anti-CD40 blocking antibody (clone W17212H) (Biolegend, San Diego, CA, USA) at 1 μg/mL was included with T cells and infected macrophages for detection of CD40L expression on T cells, as described (Gail et al., 2024, 2023). For scRNAseq experiments, anti-CD40 mAb blockade was *not* performed. Instead, PE-Dazzle-594 anti-CD40L mAb (clone 24-31) (Biolegend) was added to identify CD40L that may become internalized after ligation with CD40, avoiding blockade, as described (Musvosvi et al., 2023; Gail et al., 2024). CD4^+^ T cells were co-cultured with infected macrophages in cRPMI at 37°C and 5% CO_2_ overnight. A cocktail of anti-MHC-II blocking antibodies (anti-HLA-DR, clone L243, anti-HLA-DR,DP,DQ, clone Tu39, No Azide, Low Endotoxin grade) (BD Biosciences, Franklin Lakes, NJ, USA) were added (final concentration of 25 μg/mL, each) to some Mtb-infected samples prior to the addition of T cells as negative controls. Positive controls included stimulation with 1ug/mL staphylococcal enterotoxin B (SEB) (Toxin Technology, Inc., Sarasota, FL, USA) or Mtb lysate (for LTBI samples).

### Flow cytometry staining and analysis

After 16hrs, T cells were harvested with AutoMACS Rinsing Buffer (Miltenyi). For a subset of experiments, the Miltenyi IFNγ Secretion Assay was performed according to the manufacturer’s instructions, followed by viability dye Live-Dead Aqua or Violet (Invitrogen) staining, as described (Gail et al., 2024). Immunostaining was performed with fluorescently-labeled antibodies and FcR block (Biolegend) diluted in AutoMACS running buffer. After 20 min at 4°C, cells were washed and fixed in 1% PFA in PBS for 1 hour, followed by removal from the BSL-3 lab. Samples were acquired on the LSR Fortessa X-20 Cell Analyzer (BD). Fluorescent antibodies used in flow cytometry and sorting experiments included: APC or BUV395-conjugated anti-human CD69 (FN50), BUV737 anti-CD3 (UCHT1), BV650 anti-CD45RA (HI100), BUV395 anti-CD4 (RPA-T4), and BB515 anti-CD25 (M-A251), from BD biosciences, and BV785 anti-CD4 (RPA-T4), PE anti-TCRβ (H57-597), FITC anti-CD19 (HIB19), and PE-Dazzle-594 anti-CD40L (24-31) from Biolegend.

### Flow sorting of activated T cells for scRNAseq

Non-adherent cells were harvested from T cell-infected MDM co-culture, and immunostaining was performed, as described (Gail et al., 2024). After for 20 min at 4°C, cells were passed through 40μm filter top tubes (Corning). Approximately 2 min prior to sorting each sample, 7-AAD viability dye (Invitrogen) was added to each sample. Samples were sorted using the “purity” setting, gating on 7-AAD^lo^ CD4^+^ singles cells, and co-expression of CD69 and either CD40L or CD25. Sorting was performed on a Sony MA900 Multi-Application Cell Sorter (Sony Biotechnology, San Jose, CA, USA). Fluorescent antibodies used in flow sorting experiments include FITC anti-human CD4 (clone RPA-T4), PE anti-CD25 (clone BC96), PE-Dazzle-594 anti-CD40L (clone 24-31), BV605 anti-CD8α (clone RPA-T6), and APC anti-CD69 (clone FN50) from Biolegend.

### Exogenous antigen stimulation of CD4^+^ T cells

Exogenous antigens were added to some samples of Mtb-infected MDMs for co-culture with autologous CD4^+^ T cells. H37Rv whole cell lysate (BEI Resources, NR-14822) was added to Mtb-infected MDMs at a final concentration of 10 μg/mL for 24h. Lysate was washed out with cRPMI prior to the addition of CD4^+^ T cells. For evaluation of T cell responses to MTB300 peptides (Arlehamn et al., 2013), MTB300 was diluted to 1 μg/mL in cRPMI, then added to Mtb-infected MDMs ∼ 10 minutes prior to adding CD4^+^ T cells in co-culture. For each participant, PBMCs were also separately stimulated with either MTB300 or viral and vaccine “control” peptide megapools to identify TCR clonotypes linked to antigen specific T cell responses. For Mtb peptide stims, 1μg/mL of the MTB300 peptide megapool was added to 10×10^6^ PBMCs in cRPMI. For “control” peptide stimulations, another 10×10^6^ PBMCs were exposed to a final concentration of 1μg/mL each of Cytomegalovirus (CMV), Epstein-Barr virus (EBV), Pertussis (P), and Tetanus toxoid (TT) peptide megapools(Dan et al., 2016; Arlehamn et al., 2016; Pro et al., 2015; Antunes et al., 2017), and 1ug/mL of the SARS-CoV-2 USA-WA1/2020 Coronavirus Spike (S) protein overlapping peptides (BEI resources NR-52402). Peptides were added to PBMCs in cRPMI together with anti-CD28/CD49d costimulatory mAbs (BD Biosciences) and anti-CD40 blocking mAb (biolegend) for 16-18hrs. Following stimulation, CD4 positive immunomagnetic selection was performed according to manufacturer’s instructions using human anti-CD4 microbeads (Miltenyi Biotec), followed by flow sorting of the activated CD4^+^ T cells for downstream scTCRseq.

### Sample processing for scRNAseq

Sequencing of mRNA from single T cells was performed using the 10x Genomics Chromium Next GEM Single Cell 5’ V2 platform (10X Genomics, Pleasanton, CA, USA). Activated CD4^+^ were resuspended in 0.04% BSA in PBS and loaded onto the 10X Chromium Controller. RNA was reverse-transcribed, and cDNA was isolated and PCR-amplified according to the 10X Genomics NextGEM 5’ V2 scRNAseq User Guide (Rev D). Quality control and quantification of cDNA and final libraries was performed on either the Agilent fragment analyzer or Agilent 2100 Bioanalyzer (Agilent technologies, Santa Clara, CA, USA). V(D)J amplification and library preparation, and gene expression library construction were performed using 10X Genomics kits according to the User Guide. Paired-end sequencing was performed by the Cleveland Clinic Lerner Research Institute Genomics Core (Cleveland, OH, USA) using an Illumina NovaSeq 6000 sequencer (Illumina, San Diego, CA, USA) according to the 10X Genomics User Guide. Demultiplexed FASTQ files were mapped to the human genome and TCR sequences, simultaneously, using Cell Ranger Multi V7.1 (10X Genomics) on the 10X Cloud server and the GRCh38.p14 reference genome.

### scRNAseq analysis

Filtered_contig_annotations.csv and clonotypes.csv files from Cell Ranger containing productive, full-length TCR sequences for each sample were used for TCR clonotype analysis. Filtered feature barcode matrices (mRNA data) and contig annotations (CDR3α and β sequences) were linked to individual cell barcodes. Gene expression and TCR annotations were generated for each cell for simultaneous TCR and transcriptomics analysis after quality control and integration were performed using the R-based Seurat package (v 5.1.0) (Satija et al., 2015) with seed 123. High-quality cells and features were selected based on the following parameters: transcript detection in at least 3 cells, 200<nFeature<4,000, and percentage of mitochondrial RNA is less than 20% per cell. TCR information was added to Seurat objects using the scRepertoire package (Borcherding et al., 2020). TCR clonotypes with identical CDR3α/β sequences, present in more than 2 cells (based on unique barcodes) were considered “expanded”. Clonotypes with only an individual CDR3α or CDR3β chain were classified as “non-expanded”. After merging the samples, SCTransform (Hafemeister and Satija, 2019) was used for normalization, the percent mitochondrial and ribosomal counts, and cell-cycle gene scores were regressed out, and TCR genes were removed from the variable feature list using the quietTCRgenes function from the Trex package (Borcherding et al., 2024). Principal component analysis (RunPCA function) was followed by data integration and batch correction with the Harmony algorithm (Korsunsky et al., 2019). A UMAP (via RunUMAP) was calculated using the first 30 dimensions of the Harmony embedding with default parameters except the number of neighbors was set to 50 in addition to the FindNeighbors default parameters. Clustering was done on this shared-nearest neighbor graph using the FindClusters default parameters except the resolution was set to 0.3 using the Louvain algorithm with multilevel refinement. From 19 clusters obtained (from 0 to 18) (**Fig. S4A**), clusters 17 (27 cells) and 18 (25 cells) were removed due to low cell numbers, and cluster 16 (830 cells) due to monocytic contamination. A total of 157,462 cells split into 16 clusters were left for the downstream analysis.

### Clonal TCR analysis

The total set of TCR sequences from LTBI vs. non-LTBI participants were compared for TCR clonality/diversity using entropy indices (Shannon and Inverse Simpson) (Stewart et al., 1997). Unique αβTCR sequences (clonotypes) present in ≥ 2 copies were used to compare the responses to Mtb-infected macrophages ± MTB300 peptides, or infected macrophages ± lysate. A combined list of the unique (and total) TCR clonotypes identified in response to Mtb-infected macrophages + MTB300 peptides was generated from all experiments which evaluated a response pair (infected macrophages ± MTB300). The list of TCRs was compared to the combined list of TCRs from responses to infected macrophages only, to determine the percent recognizing infected macrophages (or MTB300 peptides only). The same approach was taken for paired analysis of infected macrophages ± lysate.

### GLIPH2 TCR analysis

A list of all HLA-II alleles for each participant, and a combined list of all CDR3β, CDR3α, Vβ, and Jβ genes, and number of clones, for expanded TCRs (≥ 2 copies) from each participant and experimental condition was generated. In cases where two CDR3β sequences were identified in a cell, they were separated and each linked to the same CDR3α, V, and J genes for compatibility with GLIPH2. HLA typing was performed by the University Hospitals of Cleveland Histocompatibility and Immunogenetics Laboratory (Cleveland, OH, USA). GLIPH2 analysis was performed as described (Huang et al., 2020) using BLOSUM62 restriction for interchangeable amino acids. Using the jvenn (Bardou et al., 2014) Venn diagram builder, GLIPH2 groups containing TCRs linked to a response to control peptides were separated from analysis. TCR sequences within GLIPH2 groups were cross-referenced with TCRs annotated in the immune epitope database (IEDB), and we removed entire GLIPH2 groups that contained any CDR3βs that contained ≥ 97% homology with TCRs previously annotated as specific for viral antigens in other studies.

To increase robustness of GLIPH2 groups, a TCR dataset containing >100 participants from 3 published studies, from Musvosvi, et al. (Musvosvi et al., 2023), and TCRs from Cleveland participants was added, followed by re-analysis using GLIPH2. In addition to the statistical criteria above, only GLIPH2 groups containing TCRs from ≥ 3 participants and an HLA association score < 0.1 were evaluated. Sequence logo plots of the CDR3β and α chains were generated using WebLogo3 (Crooks et al., 2004). GLIPH2 groups with consistent CDR3α homology (differing by ≤ 2 amino acids) were identified for transcriptomics mapping.

### Bulk TCRβ deep sequencing

Unstimulated CD4^+^ T cells were isolated by immunomagnetic positive selection from 10×10^6^ PBMCs from each participant using human anti-CD4 microbeads (Miltenyi Biotec). Genomic DNA was isolated using the QiaAMP DNA Mini Kit (Qiagen, Hilden, Germany) according to the manufacturer’s instructions. Ultra-deep level TCRβ sequencing was performed by Adaptive Biotechnologies (Seattle, WA, USA). Data was downloaded from the ImmunoSeq Analyzer (Adaptive) and used to estimate the natural circulating frequencies of TCRs. TCRβ sequences were identified in the scTCRseq analysis of CD4^+^ T cell responses to Mtb-infected macrophages, cross-referenced to CDR3β sequences in bulk TCRβ deep sequencing data from PBMCs, which were enumerated using Microsoft Excel and graphed in GraphPad Prism.

### Single-cell transcriptomics analysis

Gene expression visualization was performed using the Nebulosa package (v 1.16.0) (Alquicira-Hernandez and Powell, 2021). To identify cluster-defining genes, the FindConservedMarkers function was run on the integrated dataset with following parameters: min.pct = 0.25 and logfc.threshold = 0.25; the results were arranged according to average log_2_FC and the top 5 genes for each cluster were visualized in a dotplot in Fig. 5 using the scCustomize package (v 2.1.2). Single-cell trajectory analysis and projection was done using the Monocle3 package (Cao et al., 2019). The list of DEGs for each cluster was determined from the integrated dataset and running FindAllMarkers function with the following parameters: min.pct = 0.25, logfc.threshold = 0, test.use = "MAST", recorrect_umi = FALSE. Genes upregulated in selected clusters were visualized with the dittoSeq package (v 1.19.0) (Bunis et al., 2020). Reactome pathway gene set enrichment analysis (Milacic et al., 2023) was performed using the clusterProfiler package (v 4.14.4) (Xu et al., 2024). Modeling of intercellular communication (i.e., ligand activity analysis, target gene prediction, and ligand-receptor interactions) was performed using the nichenetr package (v 2.2.0) (Browaeys et al., 2020).

### Cloning of TCRs

TCR sequence transfection methods were adapted from Dezfulian et al (Dezfulian et al., 2023). Nucleotide sequences for selected TCRs were synthesized (Twist Biosciences, Boston, MA, USA) and inserted into pDONR221 Entry Vector (Invitrogen). All TCRs were engineered as hybrid TCRs, in which the entire human α and β V(D)J regions were preserved, and these nucleotides were grafted onto mutant mouse α and β constant regions, respectively, and cloned into the pDONR221 Entry vector as described (Dezfulian et al., 2023). Mutant mouse constant regions were used to increase TCR abundance by removing a degron within the TCRa chain, to enhance TCR association with CD3 for more efficient T-cell signaling, and to improve specific pairing and identification of the cloned TCRs. The codon-optimized nucleotide sequence for hybrid αβTCR sequences were synthesized (Twist Biosciences) as a single polyprotein separated by a 2A self-cleaving peptides sequence (TCRβ-P2A-TCRα) and subsequently cloned into the pDONR22 entry vector and then transferred into the pHAGE-EF1:DEST-PGK:CD4 destination vector provided by Dr. Mohammad Haj Dezfullian, using LR Clonase (Invitrogen) according to Gateway cloning methodology (Invitrogen) and as described (Dezfulian et al., 2023), and used to transduce 2T1R competent cells (Invitrogen). Selection of target cells was performed using ampicillin (100 μg/mL) (Gibco).

Lentivirus was produced through the co-transfection of packing plasmids (Tat, Rev, Gag-Pol, and VSV-G) (Addgene, Watertown, MA, USA) and destination vector into the HEK293T cell line (Takara Biosciences, Kusatu, Shiga, Japan), using jetPRIME® (Polyplus, Illkirch, France). Lentiviral supernatants were collected 48h after HEK293T transfection, filtered with a 0.45μm filter (Millipore), and concentrated using 100k Macrosep Advance Centrifugal Devices (Pall Corp., Fort Washington, NY, YSA).

Transduction using lentiviral supernatants was performed in the SKW-3 cell line (Schrieb Leibniz-Institut DSMZ, Germany). Concentrated lentivirus was added to SKW3 cells suspended in cRPMI at 1×10^6^ cells/mL, then incubated for 72 hours at 37°C in 5% CO2. Transduced cells were washed, then positively selected using immunomagnetic selection using biotinylated or PE-conjugated anti-mouse TCR-β, followed by anti-PE microbeads (Miltenyi Biotec) according to the manufacturer’s instructions. Antigen screening was performed either pre- or post-selection using the SKW-3 cell line transduced with the TCR of interest.

### Generating donor-specific APC cell lines

Autologous EBV-transformed B cell lines were prepared according to the ATCC Lymphocyte transformation protocol (www.atcc.org). Briefly, irradiated MRC-5 feeder cells (ATCC-55X) (ATCC, Manassas, VA, USA) cells were plated in flasks. Three days later, B cells were isolated from PBMCs by immunomagnetic negative selection using human anti-CD3 and anti-CD14 microbeads (Miltenyi Biotec). The negative fraction was co-incubated with the MRC-5 feeder cells and transformed by adding human gammaherpesvirus (HHV-4; ATCC-VR 1492). After 14-21 days, when signs of successful transformation are visible, cells were transferred to larger flasks, expanded in culture, and then cryopreserved for future use.

### Antigen screening of cloned TCRs

TCR-transduced SKW3 cells were expanded and then co-cultured 1:1 with donor-matched EBV-transformed B cells loaded with each of the fifteen 20-peptide “subpools” of MTB300, at final concentrations of 1μg/mL, overnight at 37°C in 5% CO_2_. DMSO (Sigma), MTB300 (1μg/mL), and CD3/CD28 Human T-activator Dynabeads (Gibco) were used as controls. Immunostaining with anti-CD4, CD19, TCRβ, and CD69 along with Live/Dead was performed, followed by fixation in 1% PFA/PBS. The portion of TCRβ+ cells expressing CD69 was enumerated by flow cytometry. Upon observing reactivity to a subpool, the assay was repeated using each of 20 individual peptides (at 1μg/mL) for the subpool, in addition to controls, to identify the peptide that led to CD69 expression.

### Statistics

Flow cytometry was analyzed using FlowJo V10 (BD Biosciences, San Diego, CA). Graphs and statistical comparisons were performed on Prism V10 (GraphPad, San Diego, CA, USA) using tests specified in figure legends. Grouped data were compared using non-parametric tests (Wilcoxon matched-pairs signed rank test). Data were tested for normality using the Shapiro-Wilk or Kolmogorov-Smirnov tests in Prism V10. If normally distributed, experiments with 2 conditions were compared using a paired or unpaired t-test with Welch’s correction. Two-tailed p values < 0.05 were considered significant. For single-cell transcriptomics analysis, per-cluster quantification of cell counts, clonal sizes, and CDR3 motifs was done using the R package Speckle (v 1.7.0) (Phipson et al., 2022). GLIPH2 groups considered for further analysis contained either > 2 unique CDR3β sequences, TCR Vβ gene homology score (p < 0.05), CDR3β length distribution score < 0.05, and a Fisher exact score (p < 0.05) for distinct CDR3β motifs from the V1 reference set of TCRs. HLA association score for GLIPH2 analysis of combined TCR datasets (one-sided p < 0.05), as reported previously (Musvosvi et al., 2023). Differential gene expression analyses were performed using the Seurat function FindAllMarkers to identify significantly expressed genes at p < 0.05.

## Data availability

Primary data will be available in the Supplemental Materials upon publication. The original code reported in this paper will be available through a GitHub link. The original sequencing data will be available in a public repository for future analyses. Other data will be provided by the corresponding author upon reasonable request upon publication.

## Acknowledgements

The following reagent was obtained through the NIH HIV Reagent Program, Division of AIDS, NIAID, NIH: Peptide Pool, Human Cytomegalovirus (HCMV) pp65 Protein, ARP-11549, contributed by DAIDS, NIAID. The following reagents were obtained through BEI Resources, NIAID, NIH: *Mycobacterium tuberculosis*, Strain H37Rv, whole cell lysate, NR-14822, Mycobacterium tuberculosis, Strain H37Rv, Mycobacterium tuberculosis strain H37Rv, NR-13648, and Peptide Array, SARS-Related Coronavirus 2 Spike (S) Glycoprotein, NR-52402. This work was supported by National Institutes of Health (NIH) grants K08 AI163407 (to S.M.C.), and research funds from University Hospital Cleveland Medical Center (to S.M.C.). The use of core facilities was supported in part by NIH Grant P30 AI036219 to Case Western Reserve University and the University of Pittsburgh Rustbelt Center for AIDS Research (CFAR). We are grateful to Professors W. Henry Boom and Clifford Harding for thoughtful scientific discussion.

## Author contributions

Conceptualization, S.M.C.; Methodology, V.S., D.P.G, S.R., V.G.S., A.K.S., A.S., M.H.D., C.L.A., S.M.C.; Investigation, V.S., D.P.G, S.R., V.G.S., S.M.C.; Resources, A.S., M.H.D., C.L.A., S.M.C.; Formal Analysis, V.S., D.P.G., S.R., S.M.C.; Writing-original draft preparation, V.S., D.P.G., S.R., S.M.C.; Writing-review and editing, V.S., D.P.G, S.R., V.G.S., M.H.D., C.L.A., S.M.C.; Visualization, V.S., D.P.G., S.R., S.M.C.; Supervision, S.M.C.; and funding acquisition, S.M.C.

## Supplemental Figures

**Supplemental Figure 1. CD69 and CD25 co-expression reveals a subset of memory CD4^+^ T cells that lacks recognition of Mtb-infected macrophages.** Related to Figure 1. **(A)** Flow cytometry plots comparing co-expression of CD69 and CD25, gated on gated on CD45RA^Lo^ CD4^+^ T cells after 16h co-culture with Mtb-infected (MOI 4-5) macrophages either alone or with addition of exogenous antigens (MTB300 or lysate) and **(B)** with and without α-MHC-II blocking antibodies. Plots are concatenated from 3 replicates from one experiment; data are representative of 10 independent experiments each with 2-3 replicates per condition. **(C)** Summary bar graphs compare mean (± SEM) co-expression of CD69 and CD25 (10 participants), and **(D)** the difference in activation when MTB300 is added to Mtb-infected macrophages for samples from 10 LTBI and 6 non-LTBI participants. * p < 0.05, ** p < 0.01.

**Supplemental Figure 2. Gating Strategy for flow-sorting activated CD4^+^ T cells.** Related to Figure 2. A representative flow plot showing the gating of memory CD4^+^ T cells activated in response to Mtb-infected macrophages (or controls). After gating on lymphocytes (SSC Area vs. FSC Area) and single cells (FSC Height vs FSC Area), live CD4^+^ 7-AAD^Lo^ T cells were identified. Co-expression of CD69 and either CD25 or CD40L was used to identify activated T cells for sorting.

**Supplemental Figure 3. Distribution of GLIPH2 groups linked to viral or vaccine control responses.** Related to Figures 3 and 4. **(A)** Bar graph of GLIPH2 groups (x-axis) from ≥ 1 Cleveland participant or **(B)** ≥ 3 participants from combined list of Cleveland and South African TCRs estimated to be viral or vaccine antigen-specific, based on IEDB annotation (red), non-LTBI participants (grey), or control peptide stimulations (stripes), rank-ordered by sum of highest TCR copy number (y-axis) per condition. **(C)** Bar graph of mean circulating frequency of Cleveland participants’ TCRβ clonotypes (symbols) for each GLIPH2 group from the combined list of TCRs after cross-referencing unstimulated PBMCs from Cleveland participants.

**Supplemental Figure 4. Cellularity and monocytic contamination as quality control metrics post-integration.** Related to Figure 5. **(A)** UMAP plot showing unbiased clustering of the CD4^+^ populations of 6 non-LTBI (6 donors) and 12 LTBI samples (7 donors) after QC, integration, and prior to monocytic contamination removal. **(B)** Bar plot depicting cell numbers in each cluster per experimental condition. **(C)** UMAP plot showing joint density estimation for plot for CD11c and CD206 transcripts in the integrated dataset. **(D)** Dot plot showing average expression and percentage of cells expressing CD11c and CD206 transcripts in each cluster. **(E)** Heatmap showing top 5 DEGs in the integrated dataset for each cluster prior to removal of low-quality clusters, e.g. clusters 16, 17, and 18.

**Supplemental Figure 5. TCRs with inconsistent CDR3a chains map to clusters with non-specific CD4^+^ T cell responses.** Related to Figure 6. **(A)** UMAP plot with split view (based on LTBI status) with mapping of estimated Mtb-specific TCRs from listed GLIPH2 groups containing inconsistent CDR3α homology in response to Mtb-infected macrophages, and **(B)** stacked bar plots showing their numbers per cluster. **(C)** UMAP plots with mapping of TCRs annotated as viral antigen-specific but with inconsistent CDR3α homology from listed GLIPH2 groups and **(D)** stacked bar plots showing their numbers per cluster. **(E)** Total copy numbers (left axis) and percentage (right axis) ([clone count / total cells per cluster] x 100) of cells per cluster mapping TCRs linked to a response to infected macrophages from listed GLIPH2 groups containing inconsistent CDR3α homology from known or estimated Mtb-specific GLIPH2 groups, and **(F)** for annotated viral antigen-specific TCRs. **(G)** Overlaid line graph and **(H)** correlation plot of percent T cells per cluster that mapped TCRs from all 37 GLIPH2 groups estimated to be Mtb-specific (blue) and viral antigen-specific (red).

**Supplemental Figure 6. Mtb-specific TCR clonotypes communicate with other memory T cells to control infection.** Related to Figure 7. **(A)** Network plots illustrating Reactome pathways with top DEGs labeled for cluster 6, 12, 14 (top row), and clusters 4, 11, 13, and 15 (bottom row). **(B)** Heatmaps showing downstream signaling genes estimated to be linked to cell-cell communication between ligands expressed by clusters 4, 11, 13, or 15 (combined) and T cells in clusters 6 (top) and 12 (bottom). The area under the precision-recall curve (AUPRC) was used to rank the ligand activity of senders on responders (left).

