## Supplemental Figures for "Human memory CD4^+^ T-cells recognize *Mycobacterium tuberculosis*-infected macrophages amid broader pathogen-specific responses"

### Supplemental Figure 1

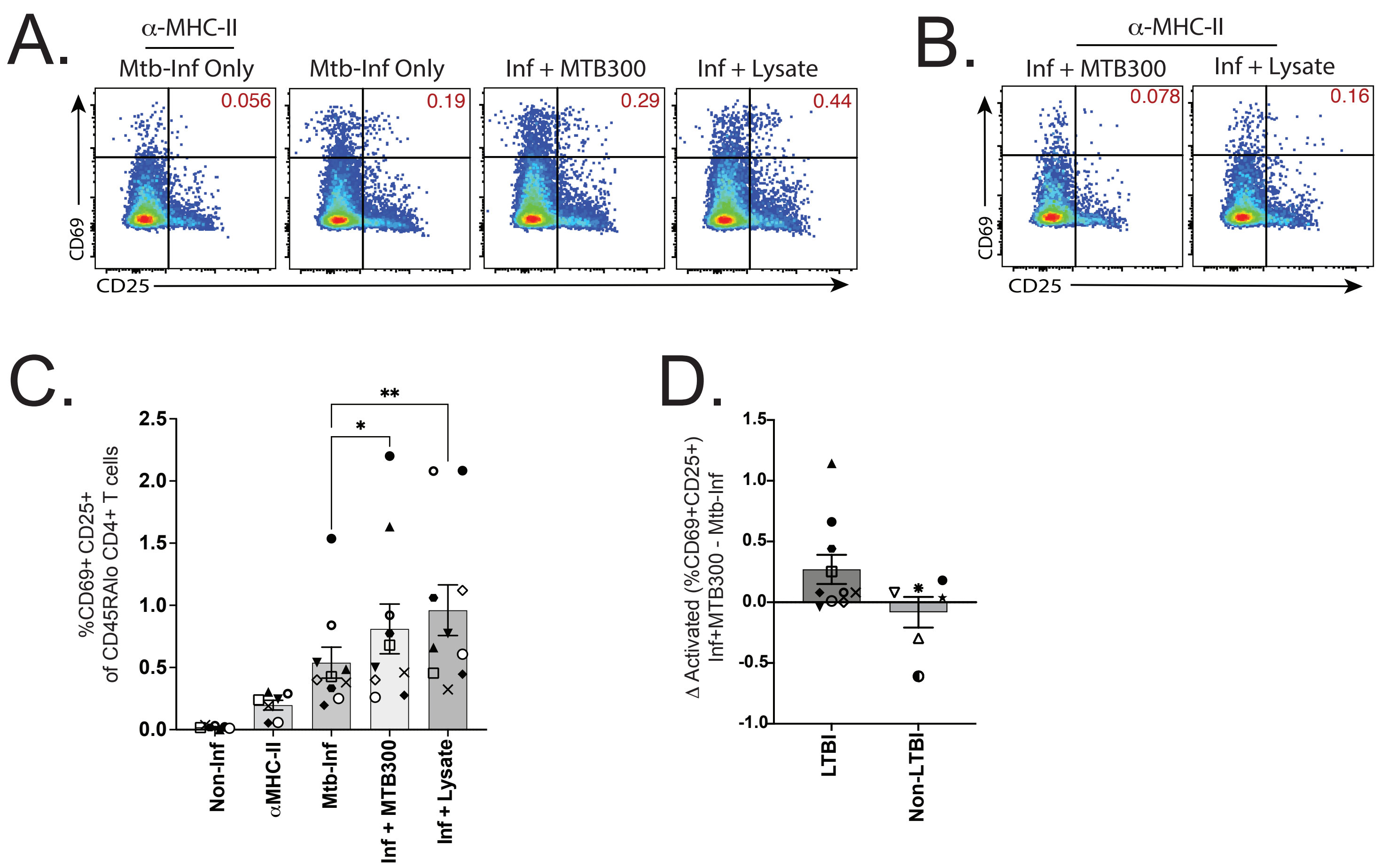

### Supplemental Figure 2

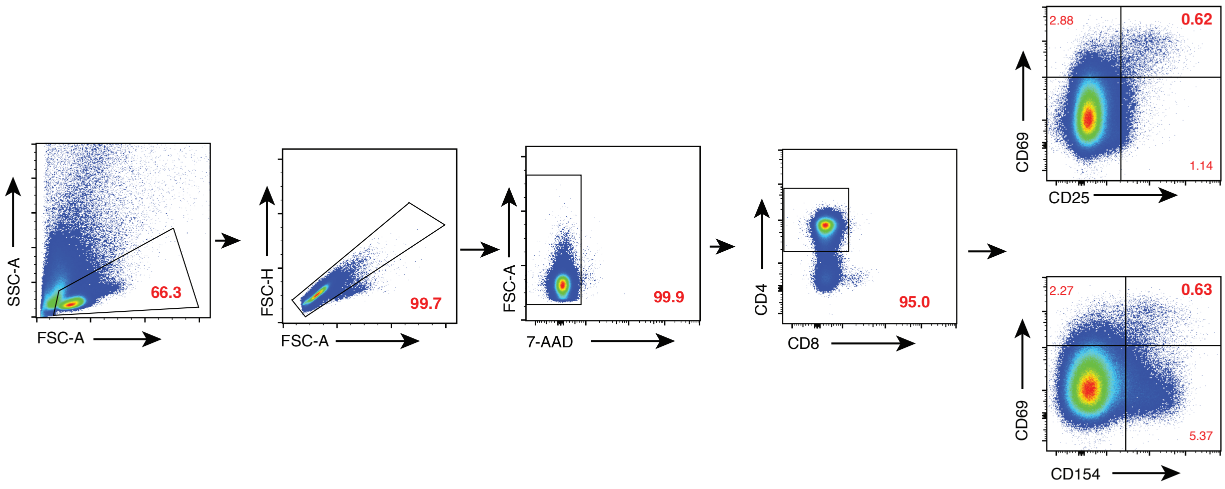

### Supplementary Figure 3

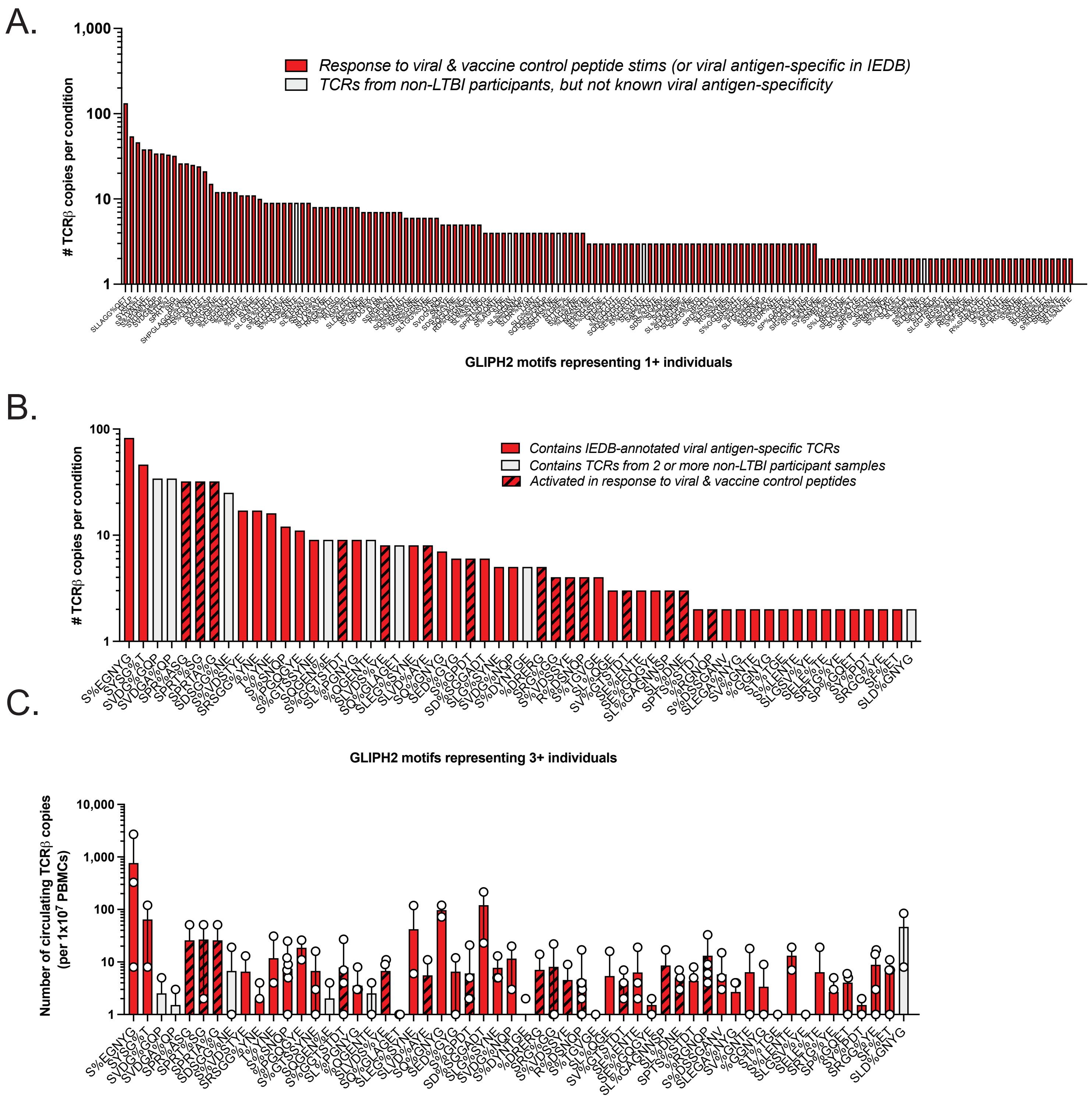

### Supplemental Figure 4

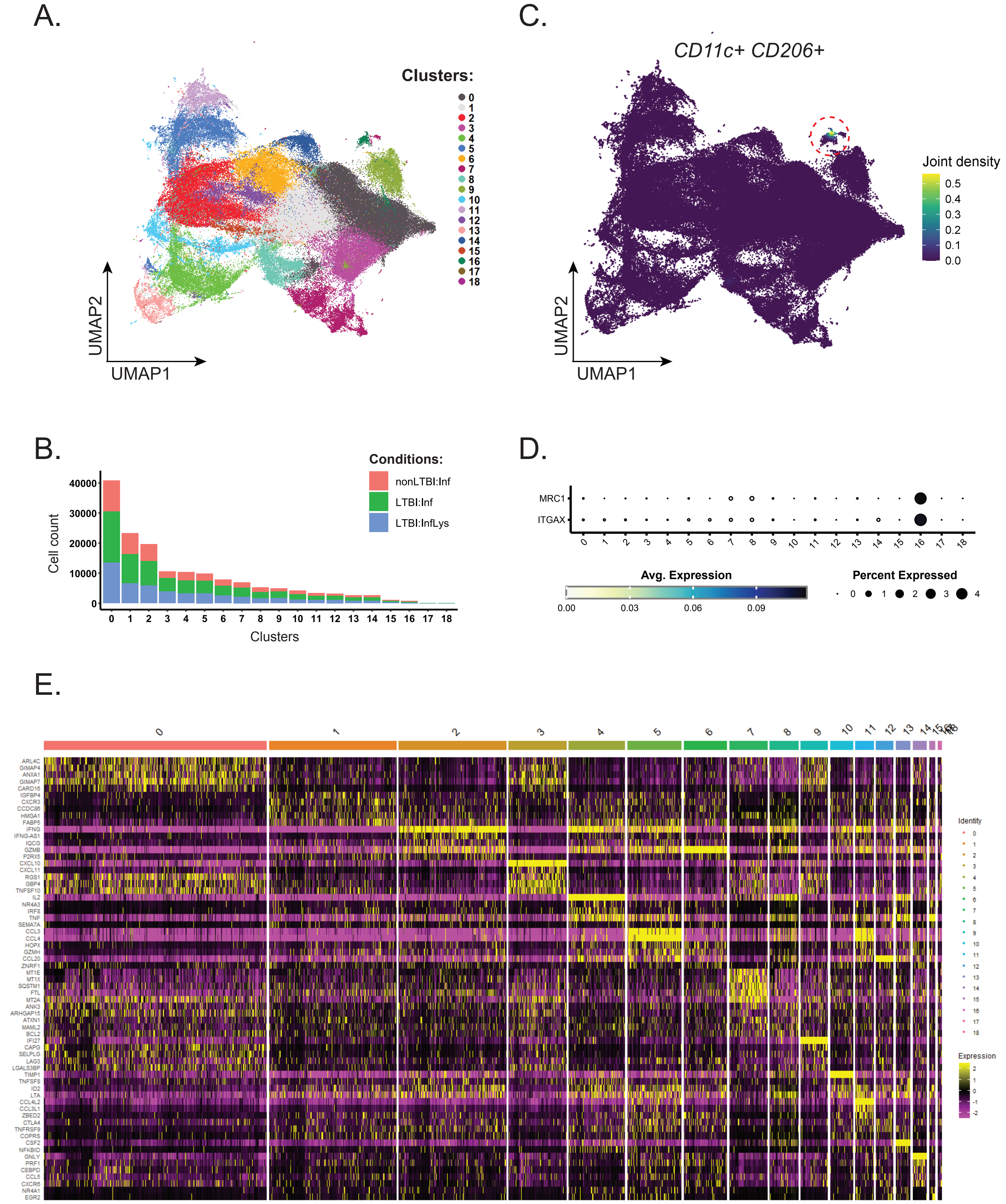

### Supplemental Figure 5

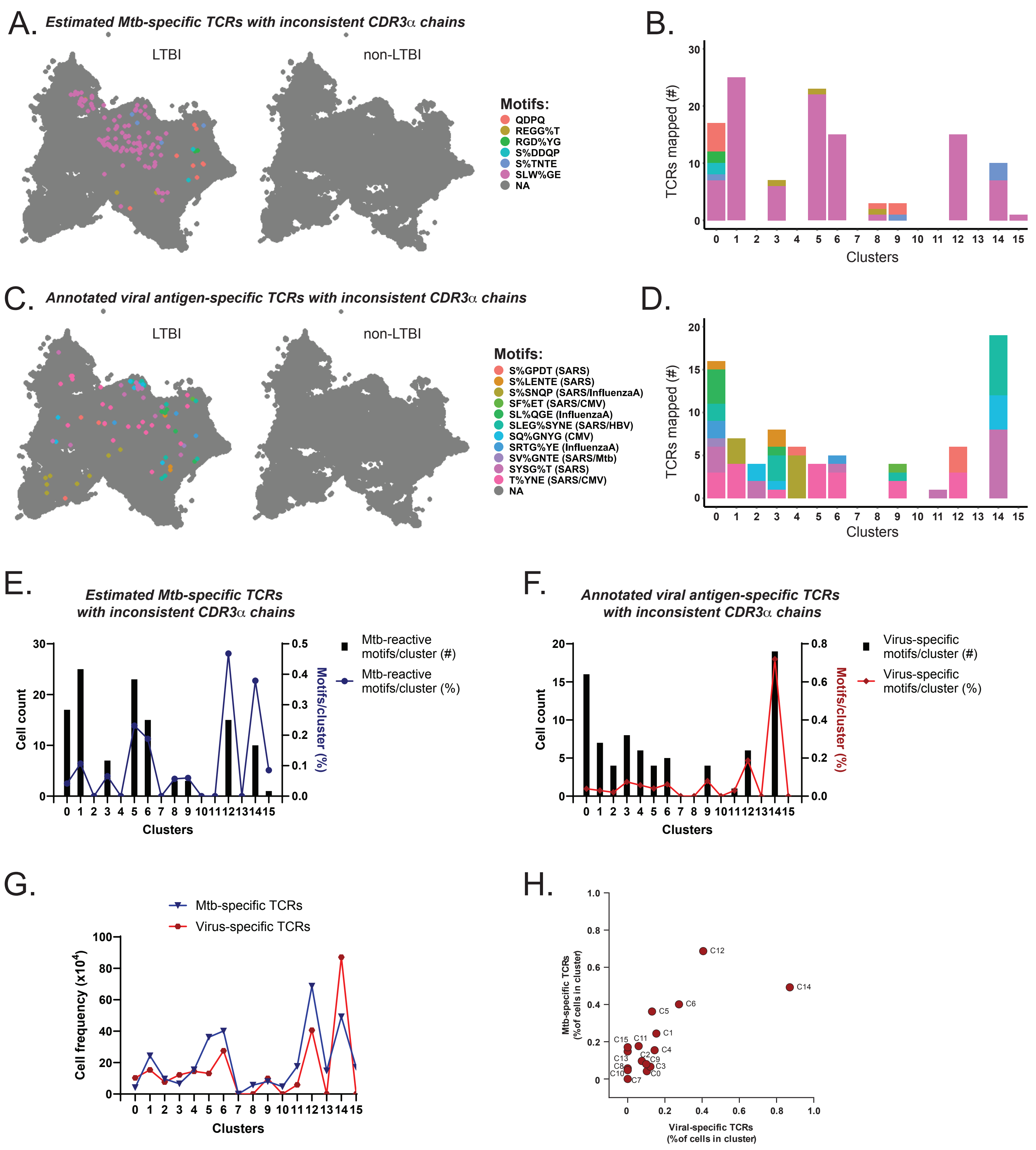

### Supplemental Figure 6

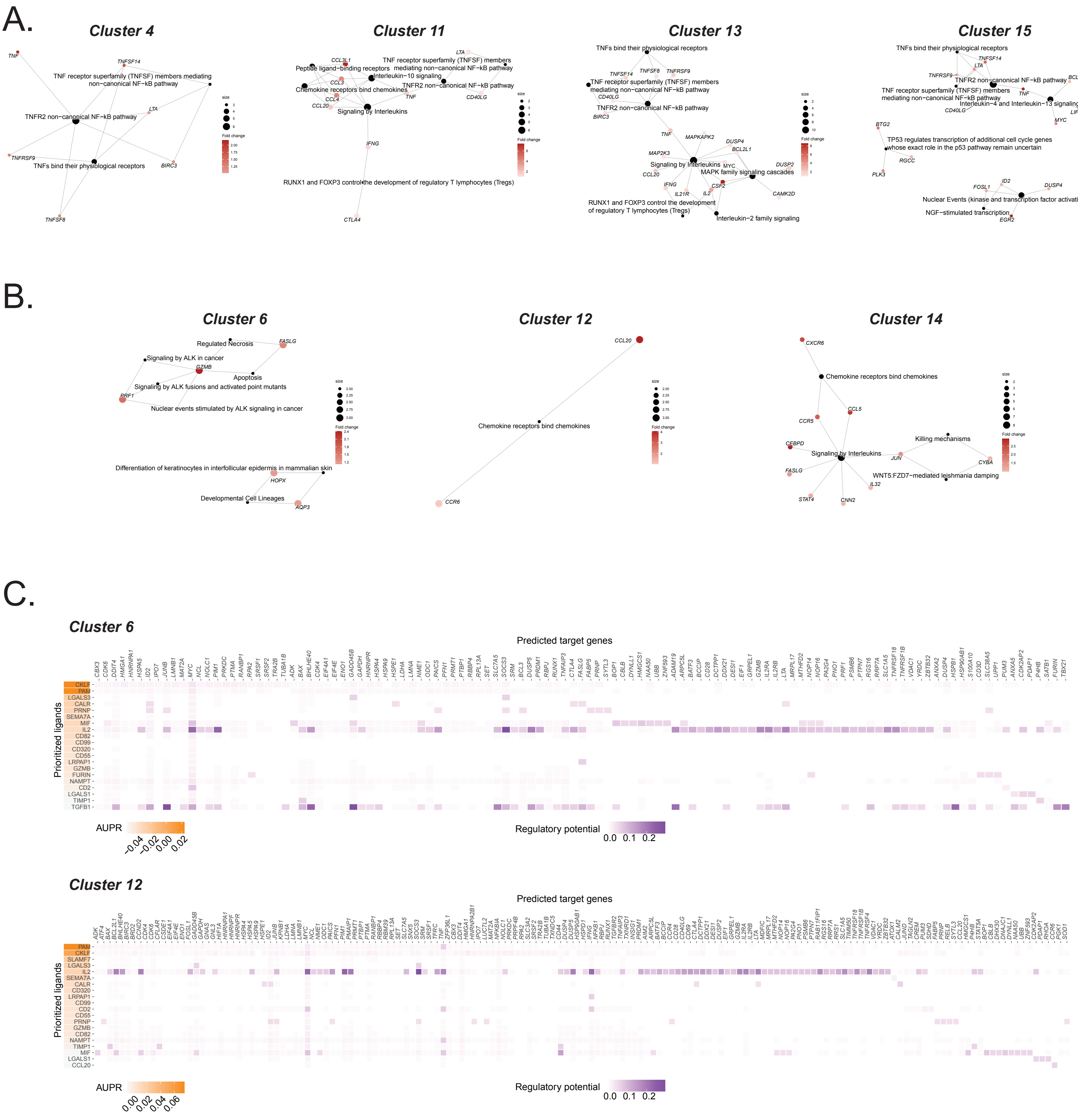
